# Magnetic DNA Origami Nanorotors

**DOI:** 10.64898/2026.03.09.710437

**Authors:** Florian Rothfischer, Lennart J. K. Weiß, Yihao Wang, Christoph Pauer, Kevin Lang, Xin Yin, Rabia Amin, Jan Lipfert, Tim Liedl, Friedrich C. Simmel, Joe Tavacoli, Aidin Lak

## Abstract

Self-assembled DNA nanostructures show great promise as functional devices, highly configurable materials, and in nanorobotics. Magnetic control can provide a powerful actuation mechanism in a broad range of contexts, since it affords a high-level of external control, it is biocompatible, and orthogonal to chemical or electrical stimuli. Here we demonstrate magnetic molecular nanoactuators by leveraging the unique site-specificity of DNA origami to assemble highly anisotropic magnetic nanocubes on high-aspect ratio DNA origami bundles. We traced and controlled 100s of our DNA origami nanorotors at the single-rotor level and demonstrated their programmable magnetic clamping and controlled rotation under uniform and rotating magnetic fields. By varying the population and inter-particle spacing of the nanocubes, magnetic torque values in the order of 10-100 pN nm are achieved at field strengths < 10 mT. Monte Carlo simulations reveal that assembly of nanocubes on DNA origami rotors leads to collective magnetic properties, with numerically estimated torque values in good agreement with the experiments. Our magnetic nanorotors offer a foundation for biocompatible nanorobotics, as well as high-throughput magnetic force and torque tweezers.

## INTRODUCTION

The application of forces and torques through magnetic manipulation at the nano– and microscales has enabled a broad range of measurement modalities and critically advanced the understanding of molecular and cellular mechanics as well as cell migration, proliferation, differentiation, and downstream signaling pathways.^1,2,11–14,3–10^ Magnetic particles provide a handle to control motion on the microscale and below via their coupling to external magnetic fields.^15^ Accordingly, they have been employed in numerous micro– and nanorobotic systems, particularly those directed towards the study of single-molecule biological systems and healthcare applications based on targeted drug delivery.^12,16–23^ Early studies mainly used ∼µm-sized magnetic beads in magnetic tweezers assays and have provided key insights into the mechanics of nucleic acids, cells, and recently proteins.^13,24–29^ However, ∼µm-sized particles provide only limited control at the molecular scale and often bind to several molecules simultaneously due to their size and multivalency. To address the bead’s multivalency issue, magnetic nanoparticles with monovalent binding ability in combination with magnetic micro-tweezers have been developed.^30,31^ However, magnetic nanoparticles with sizes < 25 nm have, inherently, a small magnetic moment m of ≈ 1-2×10^−18^ A m^2^, and are only able to generate ≈ fN forces even at few hundreds of mT.^32,33^ This force regime is orders of magnitudes too low to manipulate cellular processes, such as molecular motors and cell surface receptors that typically require 1-100 pN.^19^

Nature solves this challenge by exploiting self-assembly. Magnetotactic bacteria biomineralize a chain of iron oxide nanoparticles within their magnetosome organelles, thereby increasing their net magnetic moment by 20-fold to navigate their microenvironment following the Earth’s magnetic field.^34,35^ Such architectures can be mimicked by patterning nanoparticles on an external template and have been achieved by utilizing lithographical molding techniques, colloidal chemistry and soft synthetic templates.^16,17,44,36–43^ However, control over the number, patterning, and orientation of magnetic nanoparticles using soft and polymeric templates remains a challenge, limiting in turn the ability to precisely define the properties and functionalities of the resulting assemblies.

DNA origami, where single-stranded (ss) DNA scaffolds are folded into a 3D-nanoscale shape via hybridization of a complementary set of ssDNA oligomers, holds promise for overcoming these limitations, due to its site-specific addressability.^45–47^ DNA origami can capture and organize proteins, inorganic nanoparticles, and fluorescent dyes, when tagged with complementary base sequences, with a high spatial resolution < 1 nm.^48–54^ Moreover, DNA origami frameworks are sufficiently mechanically rigid to efficiently transfer forces and torques and to actuate defined degrees of freedom.^55–60^ Recently, dynamic DNA origami nanomachines have been developed that utilize chemical and physical triggers such as strand hybridization and displacement, thermal melting, and electric fields.^55,60–66^ Salt concentration, pH, and temperature have been employed to switch DNA origami nanomachines between pre-programmed states.^67–70^ Since these variables are tightly regulated in biological settings, this class of DNA origami nanomachines do not function under biological conditions. Electric fields have been used as an efficient physical control mechanism for DNA origami arms and rotors by coupling to their inherent charges and have shown adjustable, rapid, and repeatable unidirectional motion.^60,71^ In this manner, torque values in the order of tens of pN nm, comparable to biological rotary motors, have been achieved.^58,72–74^ However, in electrically driven DNA origami, the applied electric fields act directly on the charges of the origami structure, thus offering limited degrees of motional freedom and actuation. Moreover, an electrophoretic actuation of a 400 nm origami lever requires a 200 V bias voltage, a voltage that can heat the experimental solution and damage the DNA nanostructures and any biologically probed material.^75^

In contrast to electric fields, magnetic fields offer critical advantages for biological and biomedical applications, as they are innocuous for biological tissue and only minimally attenuated. Consequently, magnetic micro– and nano-devices are attractive for remote manipulation of biological processes and *in vivo* targeted drug delivery. Additionally, endowing magnetic properties to DNA origami nanorotors decouple function from form, thus opening a biorthogonal means of actuating without introducing heat and damaging living beings. In contrast to studies, where hybrid magnetic actuators (∼3-10 µm) were made by attaching a magnetic bead to a DNA origami micro-lever,^76^

In this study, we present magnetic DNA origami nanorotors (MADONAs) that we engineer by decorating six-helix-bundles (6HB) DNA origami nanostructures with highly anisotropic cobalt and zinc-doped ferrite magnetic nanocubes (MNCs). Our ≈ 17 nm engineered MNCs can be efficiently patterned on DNA origami and are much more magnetic than iron oxide NPs.^77,78^ Here, we demonstrate a programmable torque response by changing field strengths, particle population and spacing. Through rational design, torque values of 10-100s of pN nm at single-digit mT field strengths are achieved, thus opening new opportunities in interfacing with biological processes in the relevant torque regime on the nanoscale. Our genuinely nanoscale magnetic DNA origami nanorotors will accelerate the application of nano mechanical devices to real-world scenarios.

## RESULTS AND DISCUSSION

### Magnetic DNA origami nanorotors (MADONAs) with modular design

We build magnetic nanorotors using a ≈ 415 nm-long 6HB DNA origami as scaffold. The 6HB are designed with a fluorescently-labeled tip with 42 Atto655 fluorescent dyes to track their rotational motion using total internal reflection fluorescence microscopy (TIRFM) (Figure 1 and Supporting Information (SI)). Further, the MADONAs feature a single ssDNA-biotin extension that is tethered to the measurement glass surface via a NeutrAvidin-biotin interaction (Figure 1a and 1b) that serves as a pivot point for free hemispherical rotation. Finally, the designs have multiple binding sites for MNCs that consistent of three 8 nucleotide (nt)-long poly-dA extensions that hybridize with 20 nt-long poly-dT oligos on the MNCs.

**Figure 1.**
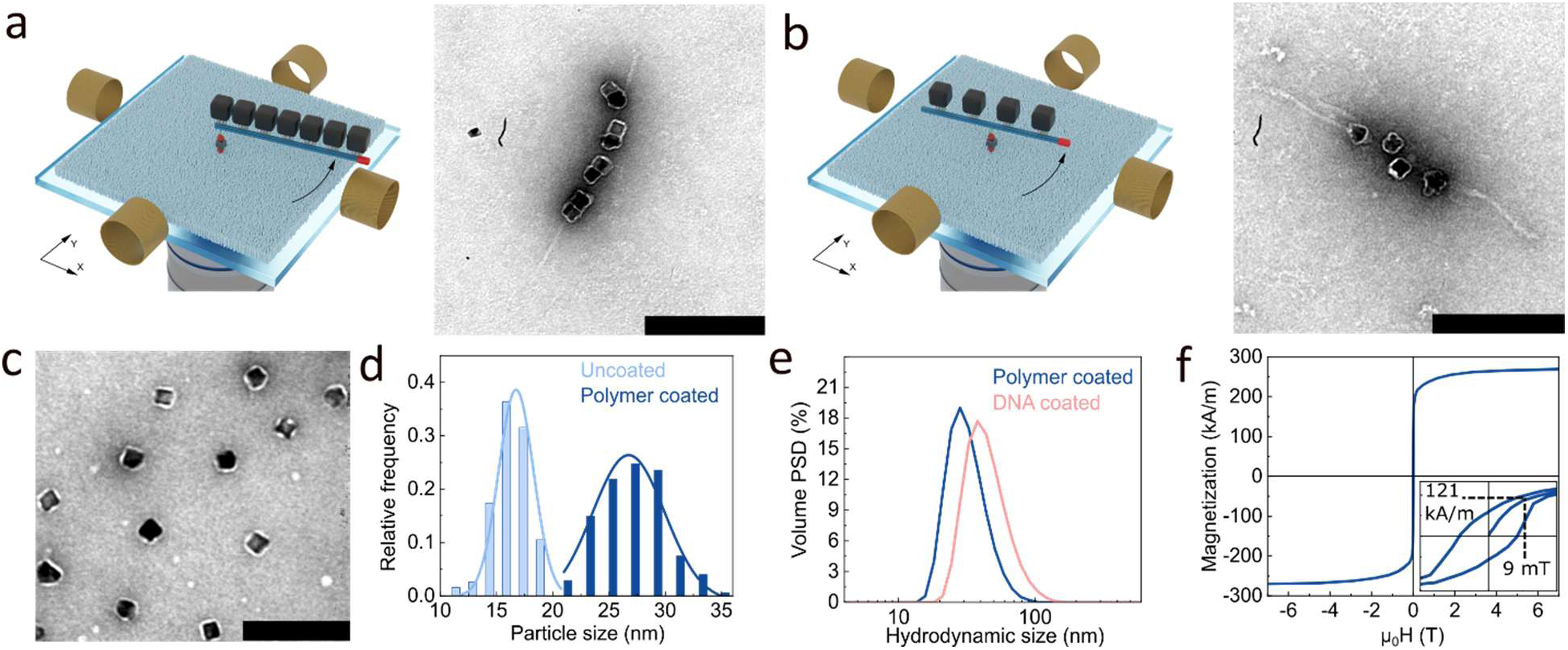
Designs of magnetic DNA origami nanorotors (MADONAs) and structural and magnetic characteristics of MNCs. (a, b) Schematic of the magnetic actuation assays at the single-rotor level (left) and negative-stain transmission electron microscopy (TEM) micrograph of a 6HB rotor (a) covered with multiple MNCs (multi-MNC design) with an end-pivot modification (right) and (b) patterned with 4 MNCs (4-MNC design) and a centered pivot point for reduced motion in z-direction. The pivot point consists of a single biotin-modification. The fluorescent tip of the 6HB is shown in red. (c) TEM image of polymer-coated MNCs. The scale bar is 100 nm. (d) Particle sizes of MNCs prior (uncoated) and after polymer coating (polymer coated) obtained from TEM analyses. (e) Volume-weighted particle hydrodynamic diameters (DLS) of polymer coated and DNA coated MNCs. (f) Magnetic hysteresis loops of colloidal MNCs. Magnetization values were converted to A m^−1^ using a density ρ = 5.28 g cm^−3^, specific to the chemical composition of our particles. At 9 mT, *M* = 121 kA m^−1^ is read from the virgin curve.

We created two different MADONA designs that vary in number, placement, and separation of their MNC-binding sites, to study how magnetic couplings between neighboring MNCs affect their magnetic response. In the first design (referred to as multi-MNC MADONA), binding sites are placed ≈ 6-8 nm from each other over a total length of ≈ 216 nm (see SI for details). The multi-MNC MADONA accommodates on average 7-8 MNCs with a particle center-to-center distance (R_c-c_) of ≈ 17 nm (Figure 1a). The continuous binding of eight MNCs along the whole structure can be seen in negative-stain TEM micrograph (Figure 1a, right panel). In the multi-MNC MADONA design, the pivot point (i.e. the center of rotation) is placed on one end of the rotor (Figure 1a). The second design (termed 4-MNC MADONA) features four defined binding sites corresponding to R_c-c_ of ≈ 64 nm and a centered pivot point. Quantitative binding of four MNCs to the second design can be seen in negative-stain TEM micrograph (Figure. 1b, right panel, Figure S4). In the 4-MNC MADONA design, we placed the pivot point at the center of the structure to control its movement along the z-axis for quantitative torque calculations (Figure 1b).

### Synthesis of colloidal MNCs with functional DNA brushes

We use custom colloidal MNCs with a dense ssDNA shell and a high magnetic moment and anisotropy to build magnetic responsive 6HB rotors.. We synthesized the MNCs following our published procedures (see the SI for protocols).^77–79^ The MNCs have the chemical composition Co_0.4_Zn_0.2_Fe_2.4_O_4_ and a mean edge length *L* of 16.7 ± 3.7 nm (N=190), as determined from the analysis of TEM images (Figure 1d, Pre-polymer coating (PC)). After encapsulating the MNCs within a thin poly (maleic anhydride-alt-1-octadecene)-PEG(3)-azide block-co-polymer shell following our protocol,^77^ the particle size *L* increases to 26.7 ± 3.2 nm (N=176), as deduced by size analysis of negatively stained TEM images (Figure 1c and d, Post-PC). After grafting 20 nt oligo(deoxythymidine) ssDNA strands on MNCs (≈ 0.12 strands/nm^2^), the particle hydrodynamic size increases from 32.6 ± 0.6 nm to 45.8 ± 1.0 nm (Figure 1e, see the SI for protocols). Magnetic hysteresis loops measured on MNCs in buffer show a saturation magnetization of *M*_s_ = 269 kA m^−1^, corresponding to an average particle magnetic moment of *m* = 1.25 × 10^−18^ A m^2^ (Figure 1f), which is roughly 8.5-fold larger than the magnetic moment of commercial 15 nm spherical iron oxide NPs.^80^ Our MNCs with an effective anisotropy constant of *K*_eff_ = 50200 J m^−3^ have an order of magnitude larger anisotropy constant than typical iron oxide NPs,^81^ and also significantly larger than 1 µm MyOne beads (details in the SI, Figure S1).^82,83^

### Multi-MNC MADONAs can be clamped at single-digit mT magnetic fields and rotated up to few Hz synchronously

We demonstrate magnetic actuation and control of our nanorotors by attaching their pivot points to a flow cell surface above a total internal reflection fluorescence microscope (TIRFM) that is equipped with two pairs of pseudo-Helmholtz coils (Figure 1a,b).^60^ The Helmholtz coils enable full field control in the (*x,y*)-plane, including the application of static or rotating magnetic fields (RMFs) of the form *B(t)* = *B*_x_(cos*(2π ft))* – *B*_y_(sin*(2π ft))*, with *B*_x_ = *B*_y_ the field strength, *f* the field frequency, and *t* time. In a first set of experiments, we attached the 6HB DNA origami with multi-particle binding sites to the glass substrate and incubated them with an excess of MNCs in the measurement channel. After washing out unbound MNCs, uniform magnetic fields are applied and the rotors’ angular position is recorded (Figure 2a and Figure 2b). At field amplitudes above 2 mT, we can clearly see that the multi-MNC MADONAs are magnetically restricted in their diffusive motion at the single-rotor level (Figure 2a). As the field strength is increased, the angular fluctuations are further restricted and we quantify the magnetic trap stiffnesses from the magnitude of angular fluctuations (Figure S4). The MADONAs return to free Brownian fluctuations upon field removal.

**Figure 2.**
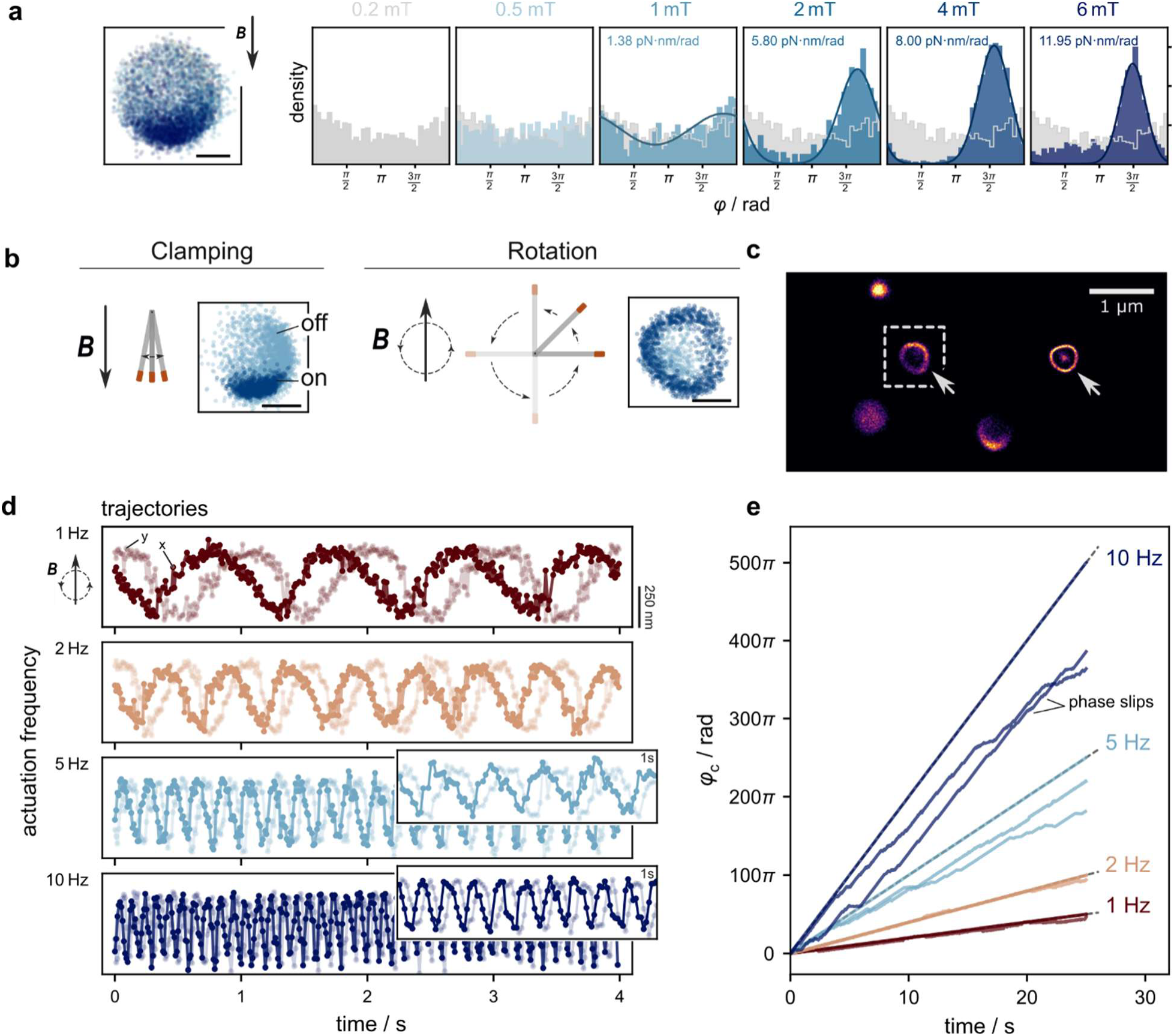
Multi-MNC MADONAs can be controlled by static and rotating magnetic fields. (a) Tracked positions of a multi-MNC MADONA rotor and histograms of the tracked rotation angle during magnetic clamping experiments at different magnetic field strengths. See also Figure S4. (b) Clamping and rotation actuation modes and their effects on rotor motion. Light blue shows the localizations during diffusion and dark blue upon magnetic actuation. Scale bar is 250 nm. (c) Localization heatmap in false colors of a TIRFM-RMF experiment. Arrows indicate functional rotors. The dashed box corresponds to rotors that follow the RMF synchronously with its rotation trajectories shown in panel (e). (d) Particle trajectories in Cartesian coordinates (scale 250 nm) of an exemplary rotor during actuation at 2.7 mT and 1, 2, 5 and 10 Hz rotating field frequencies. The insets show 1 s zoom-ins. (e) Particle trajectories shown as cumulative angle for three different rotors when driven by RFMs with the rotation frequencies indicated in the figure legend at 2.7 mT.

After clamping the multi-MNC MADONAs along a single axis successfully, we investigated their rotational dynamics under in-plane RMFs. Looking at the localization heatmap (Figure 2c) recorded under the RMFs, we observed a population of MADONAs that follow the RMF and make a full rotation circle (marked with arrows). We traced the magnetic rotations of 100s of MADONAs within a single field of view for different actuation frequencies (*x, y*-locations of a single rotor in Figure 2d, cumulative angles for three rotors in Fig. 2e). While some of the multi-MNC MADONAs synchronously follow the RMF up to 10 Hz (Figure 2e, ideal trajectories shown as dashed lines), others experience phase slips and only incompletely follow the external field, in particular at higher rotation speeds. In non-Brownian systems these so-called break down points are where the exerted magnetic torque, *τ* = *m*_eff_ *B*, matches the viscous drag and restoring torques from surface-origami interactions (see the SI for detailed discussion). Rotors with lower effective moments cannot sustain co-rotation with the RMF, in particular at lower field strengths and faster rotation speeds.

In general, we observe three rotor categories: (1) Rotors that remain freely diffusing; (2) rotors that are actuated by the field but also experience phase-slips and (3) rotors that follow the field synchronously under the conditions tested. The freely diffusing rotors are presumably non-laden with MNCs while asynchronous and synchronous rotors likely span partially to fully-loaded MNC populated rotors. Indeed, TEM images of multi-MNC MADONAs support the non-quantitative binding of MNCs due to tightly-placed binding sites (Figure S5, showing TEM images of multi-MNC MADONAs with unfilled MNC binding sites). Furthermore, rotors laden with the same number of MNCs can have different levels magnetization depending on the initial binding orientation of the MNCs that may/may not frustrate the system from reaching its lowest energy state (i.e. the highest net magnetization where the permanent dipoles of the MNCs are mutually aligned with each other and the external field). Nevertheless, our measurements clearly demonstrate that we can magnetically actuate functionalized DNA origami rotors.

### Controlled actuation by patterning four MNCs (4-MNC MADONAs) at specific inter-particle distances

Our data for multi-MNC MADONAs suggest that placing the particle binding sites close to each other leads to large variations in rotor properties, dynamics, presumably due to steric hinderance between adjacent MNCs, leading to incomplete binding (Figure S5). Accordingly, we next focused on 4-MNC MADONA design, with fewer attachment sites spaced further apart, that offers more consistent binding of 4 MNCs per rotor (Figure S6 for TEM micrographs). The combination of external actuation and TIRFM tracing at the single-rotor level enables us to experimentally determine the effective magnetic torques by successively balancing the magnetic and viscous drag torques (either by changing the magnetic field strength or the actuation frequency for the same field of view), a feature that is not easily accessible in micron-sized torque probes.^21^ First, we assessed the rotational drag coefficient *ζ* of the rotors from their diffusive motion (Figure 3a and b) and computed mean squared angular displacements to extract their rotational diffusion coefficients. The diffusion coefficients were log-normally distributed (*µ* = 2.39, *σ* = 0.55, R^2^ = 0.94) with a most probable value of *D*_rot_ = 8 rad^2^ s^−1^, which corresponds to a most likely drag coefficient of *ζ*_rot_ = 2.7 × 10^−22^ N m s. This value is in good agreement with the bulk rotational drag coefficient of 2.4 × 10^−22^ N m s for a rod with a length of 415 nm and width of 8 nm in a medium with viscosity of 12.5 × 10^−3^ Pa s,^58^ the approximate dimensions of our construct and conditions of our experiments.

**Figure 3.**
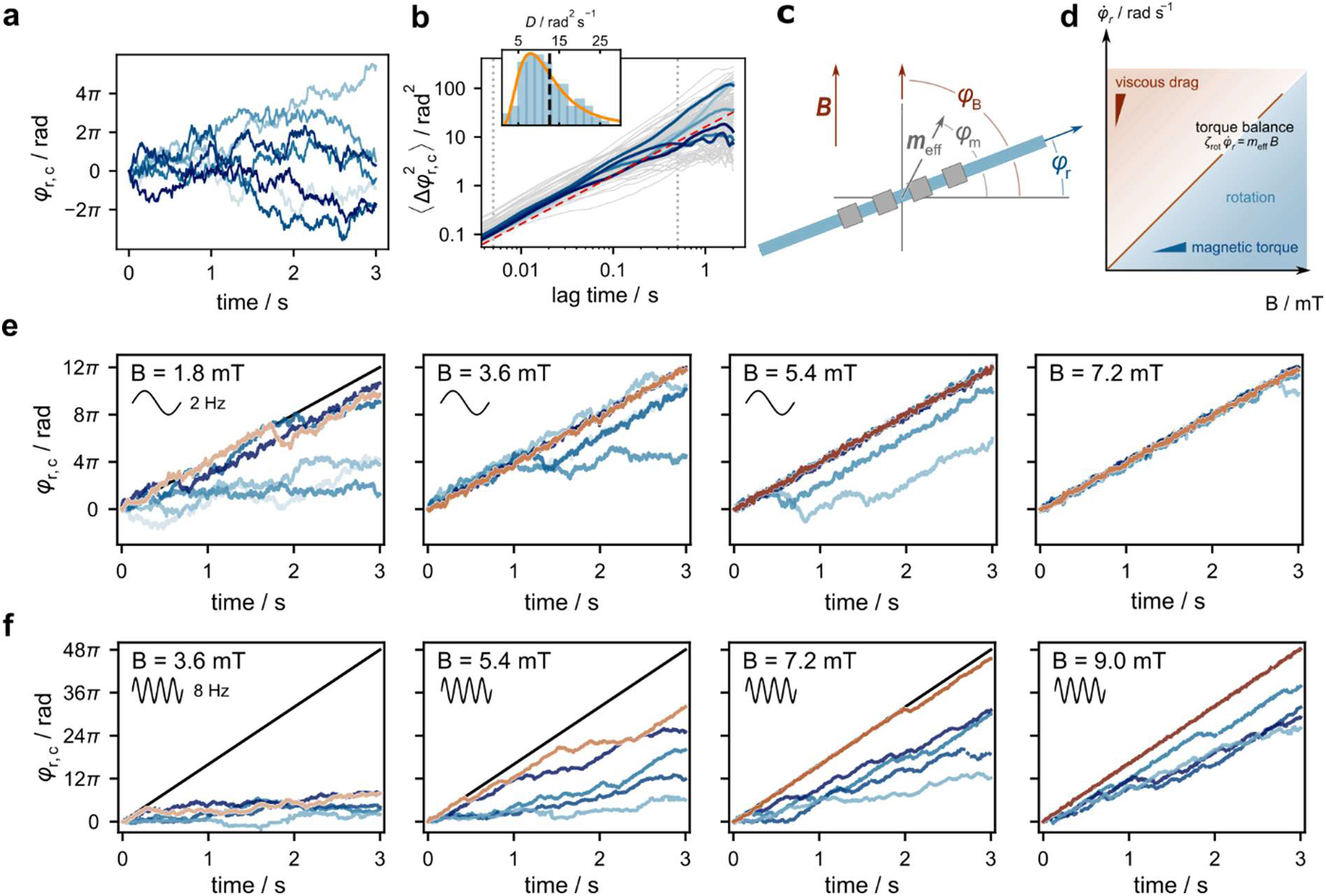
4-MNC MADONAs driven by rotating magnetic fields. (a) Diffusive rotor trajectories with the magnetic field off. (b) Mean-squared angular displacement 〈Δφ^2^〉 of the rotor ensemble during diffusion (highlighted curves correspond to trajectories in (a)). The dashed red line in the mean power spectrum. Inset: Histogram of extracted log-normally distributed (N = 128, R^2^ = 0.94) rotational diffusion coefficients, yielding an experimental mean *D*_rot_ = 12.7 rad^2^ s^−1^ and fitted most likely value *D*_rot_ = 8 rad^2^ s^−1^ (c) Coordinate system for analysis of rotor motion; *φ*_r_ is the rotor orientation, *φ*_m_ is the axis of the effective magnetic moment *m*_eff_, and *φ*_B_ is the orientation of the external magnetic field. (d) Co-rotation requires an effective magnetic moment sufficient to overcome viscous drag under external actuation. (e, f) Exemplary trajectories of nanorotors during actuation with 2 Hz and 8 Hz and different field strengths, respectively.

To understand the rotational behaviors of these rotors, we developed a rotation model (Figure 3c, details in the SI). Here, we considered a 6HB rotor that carries four MNCs with magnetic moments *m*_i_, resulting in a net magnetic moment *m*_eff_ per rotor. Upon external actuation, this effective magnetic moment experiences a torque *τ*_m_ = *m*_eff_ *B* sin(*φ*_B_ – *φ*_m_), which is dependent on its axis’ orientation *φ*_m_ relative to the external field *φ*_B_. Since the MNCs are coupled to the DNA origami, we assume a constant angle offset between *m*_eff_ and the rotor axis, *φ*_m_ = *φ*_r_ + *Δφ* (Figure 3c), which allows the rotors to co-rotate with the external field. Then, the rotor’s motion for a RMF with constant angular velocity, *φ*_B_ (t) = *ω*_B_ *t*, can be described by a torque balance in the overdamped regime, where the induced magnetic torque must compensate for the viscous drag, the local restoring torque stemming from rotor-surface interactions, and thermal fluctuations:

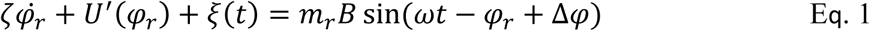

Here, *ζ* is the drag coefficient, *U*(φ_r_) is the energy landscape, and *ξ* is Gaussian white noise with 〈ξ(*t*)ξ(*t*^r^)〉 = 2ζ*k*_B_*T*δ(*t* − *t*^r^). If we neglect thermal fluctuations, the rotors are synchronous to the external drive when the angular offset between the external actuation and the rotor, θ = ω_B_*t* − φ_r_, stays constant (Figure S8-S10 for deterministic and stochastic rotor trajectories in the absence and presence of thermal fluctuations). This is possible as long as the magnetic torque compensates (at least) the viscous drag, *m*_eff_*B* ≥ ζω_B_. For smaller torques, the phase offset θ no longer locks but drifts, and the rotor lags behind the magnetic field, though net directional motion is observed. However, if we take local rotor-surface interactions (see Supplementary Notes) and noise into account, the critical limit above provides only a lower bound for the magnetic torque needed to sustain co-rotation.

We now probe the effective magnetic moment (Figure 3d) of the 4MNC-MADONAs by actuating rotor ensembles at fixed angular speed and with increasing field strength (Figure 3e for 2 Hz and Figure 3g for 8 Hz). Our TIRFM-RMF assays provide several insights into the rotors dynamics: the fraction of rotors that rotate synchronously with the RMF increases with field strength and decreases with frequency (Figure 3e and 3g). This concurs with the torque balance described in equation (1) and its magnetic moment and rotational frequency dependence. Orange-colored trajectories (Figure 3e and 3g) belong to a rotor that follows the RMF synchronously. Blue trajectories are from those rotors that are driven with the field, yet at lower angular speed, presumably due to having low *m*_eff_ and/or experiencing rough surface energy landscape. Further elaboration on quantifying magnetic torques from frequency-resolved drop-out analyses by setting the so-called phase-locking value (PLV) criterion can be found in the SI (Figure S11-S15 for rotor trajectories at different field strengths and frequencies).

### 4-MNC MADONAs exert 10s of pN nm magnetic torques at single-digit mT magnetic fields: Clamping experiments and Monte Carlo simulations

Next, we developed a 2D dynamic particle-coupling model for our ferrimagnetic MNCs (Figure 4a) to better understand how the magnetization *m* of individual MNCs interacts to produce *m*_eff_ and to provide a microscopic understanding of the clamping experiments on 4-MNC MADONAs (Figure S16 and S17). We used Metropolis Monte Carlo (MMC) algorithm to minimize the total free magnetic energy by considering the Zeeman and magnetic anisotropy energies, and dipole-dipole interactions (details can be found in the SI, Figure S18). For these simulations, we focused on 4-MNC MADONAs, as this design offers at most binding of 4 MNCs to the 6HB (see Figure S6 for TEM micrographs). We assumed that the bound MNCs can rotate physically up to a maximum of 90 ° to align with the field (Figure 4a). From the MMC simulation results (average of 15 MMC runs), we computed the rotor’s *m*_eff_ at 9 mT field strength by the vector sum of magnetic moments of four MNCs (Figure 4b). The MMC results show that the simulated *m*_eff_ of 4-MNC MADONAs is larger than the *m* of a single MNC in suspension (which was experimentally observed via SQUID magnetometry, Figure 1f and 4b). To check if there is collective couplings between MNCs in nanorotors, we compared the simulation data with the analytical expression of *m*_eff_ for a random cluster of *N* non-interacting MNPs, given by^84^

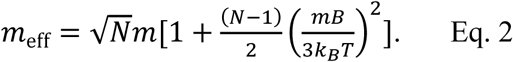

**Figure 4.**
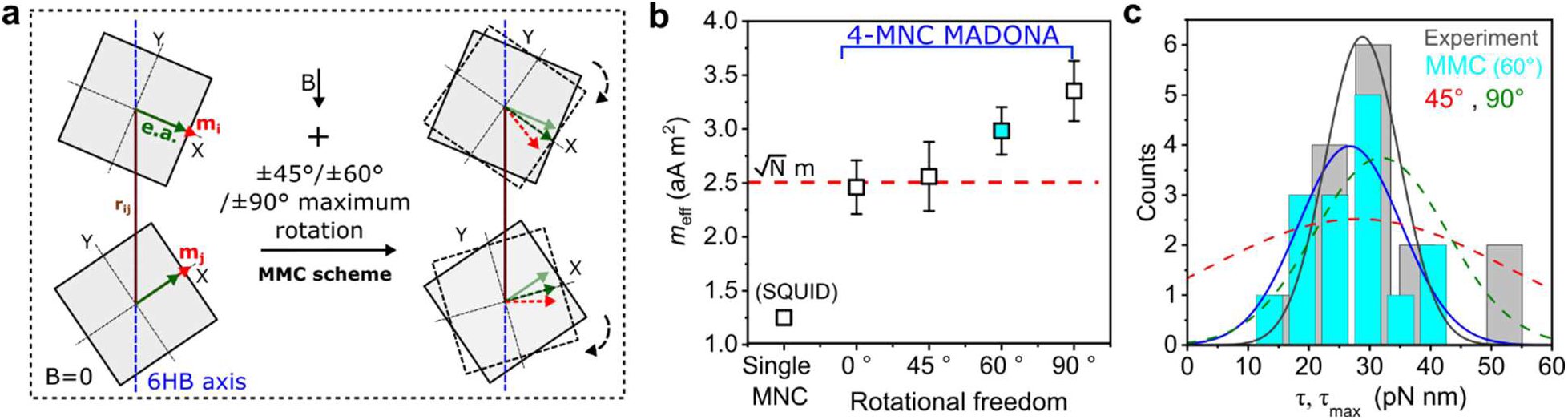
Magnetic torques of 4-MNC MADONAs from magnetic clamping and Monte Carlo simulations. (a) Physical model of magnetic interactions between two neighboring MNCs and with the external field used in Metropolis Monte Carlo (MMC) simulations. **m**_i_ and **e.a** are the particle’s *i* magnetic moment and anisotropy axis unit vectors. (b) Effective magnetic moment *m*_eff_ of 4-MNC MADONA at 9 mT field amplitudes at 0, 45, 60, and 90 ° rotational freedom obtained by from fifteen independent MMC simulations. The results are the mean values with standard error of means (S.E.M). *m*_mean_ of a single MNC measured in suspension using a superconducting quantum interference device (SQUID) magnetometer is shown for comparison. Particles parameters used in the MMC simulations are given in the SI. (c) Maximum magnetic torques obtained from the clamping assays and MMC simulations at 9 mT. The solid lines are the best Gaussian fits to the histograms. The red and green dashed lines are the Gaussian fits to the histograms of 45 ° and 90 ° rotations, with the Bhattacharyya coefficient of 0.67 and 0.90, indicating a partial and strong overlap with the experimental torque histogram, respectively. Same color coding is applied in (e) and (f).

The Eq. 2 becomes 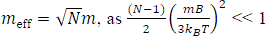 at the simulation conditions (9 mT, N = 4). We can observe that the simulated *m*_eff_ values are above 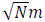 limit (dashed red line in Figure 4b) when a nominal particle rotation ≥ 60 ° is considered in the simulations. At 60 ° rotation freedom, the *m*_eff_ = 2.98 ± 0.2 aA m^2^ corresponds to a ≈ 60 % of the perfect alignment of four MNCs (i.e. *m*_eff_ = 5 aA m^2^, Figure 4b). Comparing this level of alignment with ≈ 45 % alignment of single MNCs in suspension at 9 mT (measured by SQUID magnetometry, Figure 1f), there is a 15 % alignment gain as a result of assembling of MNCs on the 6HB bundles. All together, our experimental and numerical results clearly demonstrate collective magnetic interactions between MNCs on 4-MNC MADONAs. We then computed the maximum torques *τ*_max_ by taking the simulated *m*_eff_ and compared it with *τ* values obtained from the clamping experiments (Figure S16 and S17). The best agreement between experiments and simulations was obtained at the 60 ° rotation freedom by comparing the similarity between two histograms using the Bhattacharyya coefficient (BC). The BC = 0.96 indicates a very strong overlap between experimental and simulated torque histograms (Figure 4c). Both experiments and simulations suggest that bound MNCs to the bundles are able to rotate to a certain angle in the direction of external fields, resulting in an overall stable magnetization axis of rotors.

From a biological perspective, our MADONA rotors are a highly innovative and promising class of magnetic torque nano-probes that combine three unique features. First, the torque values a single rotor can exert with only four MNCs are adequate to stall and manipulate molecular rotors such as F_0_F_1_-ATPase, requiring ≈ 40 pN nm.^85^ Second, our MADONAs can target and manipulate single molecules/receptors on membrane, here showcased by docking/rotating on a single biotin-streptavidin pivot complex. Third, due to their strong magnetic responsiveness and high anisotropy, they can be manipulated in different magnetic modes at single-digit mT field strengths, a field regime that can be achieved with a pair of low-cost permanent magnets.

## CONCLUSIONS

We present nanoscale magnetic DNA origami nanoactuators that are able to exert biologically relevant magnetic torques at single-digit mT magnetic fields. This is achieved by patterning highly magnetic and anisotropic custom MNCs on rigid DNA origami 6HBs. Our TIRFM-RMF setup enables programmable nano-rotation assays, in which the torque values were determined by balancing the drag and magnetic torques. By comparing two different rotor designs, differing in particle population and inter-particle distance, we found that quantitative binding of MNCs on the 6HB origami and thus adequate torques are achieved when the inter-binding site (particle distance) is set to more than one particle size. At this separation, there is a good tradeoff between quantitative binding and dipole-dipole interactions between MNCs. Our 4-MNC MADONAs can generate torques sufficient to stall and manipulate rotary molecular rotors such as F_1_-ATPase. Our numerical simulations back up our TIRFM-MF assays with a good agreement between experimentally and numerically torque values. Furthermore, Monte Carlo simulations show that attaching highly magnetic MNCs at 64 nm particle center-to-center distance leads to collective magnetic couplings and increased effective magnetic moment of the rotors.

Endowing DNA origami with magnetic properties by assembling bespoke MNCs on them will lead to innovative magnetic nanoactuators that can be driven into several distinct configurations within milliseconds in response to time-varying magnetic fields. Here, we show that such fast varying dynamics can be traced at the single rotor level with TIRF microscopy by tracking several dyes. In terms of understanding magnetic movements, by harnessing the binding addressability of DNA origami, our nanoplatform is an ideal platform from which to understand how complex magnetic movements can be achieved depending on particles properties and assembly of magnetic field setup.

Our magnetic nano-platform is highly adaptable and can be synthesized with high modularity and rotational degrees of freedom. Moreover, the controlled placements of nanomagnets on nanostructures allows, in principle, for the creation of a complex torque response over three degrees of rotational freedom to homogeneous and rotating magnetic fields. With this study, we set a solid foundation for working toward new class of biorthogonal nanoactuators in which the actuation is truly decoupled from the structure.

## ASSOCIATED CONTENT

Supporting Information (SI) contains: Materials and Methods, Synthesis Procedures, Supplementary Figure S1-S19. Description of rotor’s rotation dynamics. Discussion on magnetic torque determination from frequency-resolved drop-out analyses on 4-MNC MADONAs. Description of MMC simulations.

### Data Availability

Source data and all other data that support the plots within this paper and other findings of this study are available from the corresponding authors upon reasonable request.

### Code Availability

The source code of the data analysis routines and simulation files employed in this study is available from the corresponding authors upon reasonable request.

## AUTHOR INFORMATION

### Corresponding Author

* Aidin Lak

* Joe Tavacoli

### Present Addresses

#### Author Contributions

A.L. conceived the research idea. F.R. designed and folded DNA origami bundles, performed TRIFM magnetic assays, and analyzed data. L.W. performed TIRFM measurements, analyzed data and wrote the manuscript. Y.W. polymer coated and DNA labelled MNCs, performed magnetic and colloidal measurements, built a pseudo-Helmholtz coils, and analyzed particle characterization data. C.P. synthesized DNA origami, performed MNC binding to DNA origami and magnetic actuation assays. K.L. performed initial magnetic actuation assays. X.Y. folded DNA origami and performed particle binding assays and TEM imaging. R.A. synthesized MNCs. J.L. analyzed data. T.L. analyzed data and provided resources. F.S. analyzed data and provided resources. J.T. conceptualized, analyzed data and wrote the manuscript. A.L. developed MNCs and DNA labelling chemistries, performed MMC simulations, analyzed data, supervised the study, and wrote the manuscript.

## FUNDING SOURCES

German Research Foundation: LA 4923/3-1, TA1375/2-1, SFB1032 (project ID 201269156 TPA2 and TPA6)

European Research Council: ProForce (101002656)

BMBF: 6G-life initiative, ONE MUNICH Project Munich Multiscale Biofabrication

## NOTES

The authors claim no conflict of interest.

## ACKNOWLEDGMENT

We thank the group of Hendrik Dietz for providing in-house produced scaffold strands, Susanne Kempter and Kerstin Frank for assistance. This work was funded by the German Research Foundation (DFG, German Research Foundation) through LA 4923/5-1 (A.L), TA1375/2-1 (J.T), SFB1032 (project ID 201269156 TPA2 and TPA6), and Germany’s Excellence Strategy – EXC 3092/1 – BioSysteM (project no. 533751719). We also gratefully acknowledge funding by the BMBF through its 6G-life initiative, and the BMBF and the Free State of Bavaria through the ONE MUNICH Project Munich Multiscale Biofabrication and ERC Consolidator Grant “ProForce” (101002656). F.R. has personally been funded by the Konrad-Adenauer-Stiftung (KAS).

## Notes

### Competing Interest Statement

The authors have declared no competing interest.

