## Supporting Information for "Magnetic DNA Origami Nanorotors"

### CONTENTS

1. Synthesis, DNA labeling, and size fractionation of colloidal magnetic nanocubes 3
Magnetic nanoparticle synthesis 3
Functionalization of poly(maleic anhydride-alt-1-octadecene) with $NH_2$-PEG(3)-azide linker 3
MNC Polymer coating procedure 3
Particle purification and fractionation 3
Agarose gel electrophoresis to purify polymer coated NPs 4
DNA labeling of MNCs 4

2. Structural and magnetic characterization of colloidal MNCs 4
Dynamic light scattering (DLS) 4
Magnetic hysteresis loops: characterization of saturation magnetization, magnetic coercivity, and magnetic anisotropy 4
Determination of effective magnetic anisotropy constant 4
Particle size analysis 5

3. Folding and analysis of DNA origami structures 5
DNA origami folding and purification 5
Origami structure verification 6
Fluorescence microscopy sample preparation 6
Homogeneous and rotating magnetic fields 6
Analysis of single-molecule fluorescence measurements 6

4. Trap stiffness data for Multi-MNC MADONAs 9

5. TEM micrographs of rotors decorated with magnetic nanoparticles 10

6. Co-rotation as phase-locking phenomena 12

7. Drop-out analysis for rotations at different angular velocities and field strengths 18

8. Magnetic torques of 4-MNC MADONAs from frequency-resolved drop-out analyses 22
Phase-locking analysis at different frequencies 22

9. Trap stiffness data for 4-MNC MADONAs 24

10. Monte Carlo simulations of magnetic properties of MADONAs 26

11. Scaffold and staple sequences 29
$7,249$ basepair scaffold version used for the 6hb structues: 29
Staple sequences of the basic 6hb DNA origami nanostructure 32
MNC binding staples Multi-MNC 36
MNC binding staples of the 4-MNC structure 38
Modifications staples 38

References 40

### 1. SYNTHESIS, DNA LABELING, AND SIZE FRACTIONATION OF COLLOIDAL MAGNETIC NANOCUBES

#### Magnetic nanoparticle synthesis

Cobalt and zinc-doped iron oxide nanocubes (MNCs) were synthesized via thermal decomposition of iron (III) acetylacetonate, cobalt (II) acetylacetonate, and zinc (II) acetylacetonate in a mixture of dibenzyl ether, 1-octadecene, oleic acid, and sodium oleate at $290\,^{\circ}\mathrm{C}$ for 30 min as previously described [1].

#### Functionalization of poly(maleic anhydride-alt-1-octadecene) with $NH_2$-PEG(3)-azide linker

Our custom-synthesized poly(maleic anhydride-alt-1-octadecene) (PMAO) was modified with $NH_2$-PEG(3)-azide via anhydride ring opening reaction following the procedure published by us and Jin et al. [2, 3]. In a typical reaction, $338\,\mathrm{mg}$ ($0.964\,\mathrm{mmol}$ monomer units) of PMAO were added into $17\,\mathrm{of}$ dimethylformamide (DMF) in a $50\,\mathrm{mL}$ round flask and heated up to 65 °C. Next, $191.3\,\mu\mathrm{L}$ ($0.964\,\mathrm{mmol}$) of $NH_2$-PEG(3)-azide dissolved in $3\,\mathrm{mL}$ of DMF was pipetted into the reaction flask after the PMAO was completely dissolved. Afterwards, the mixture was left to react for 24 h at 65 °C. After cooling to room temperature, the crude lightly yellowish oily product was dialyzed against dichloromethane (DCM) using regenerated cellulose dialysis tubes (6 kDa cutoff size) to remove unreacted ingredients and byproducts. Finally, DCM was thoroughly removed in a rotary evaporator and the resulting polymer was dissolved in chloroform to obtain a final concentration of 35 mg (polymer)/ml and stored at 4 °C prior to usage.

#### MNC Polymer coating procedure

The polymer coating of oleic acid coated MNCs was performed and modified based on the protocol by Pellegrino et al.[4] and William W. Yu et al.[5]. Typically, $2.635\,\mathrm{mL}$ of PMAO-azide polymer (35 mg/mL) were diluted with $7.365\,\mathrm{mL}$ chloroform in a $30\,\mathrm{mL}$ glass vial and sonicated for $10\,\mathrm{min}$. Next, $2\,\mathrm{mL}$ of particle suspensions in chloroform ($1.44\,\mathrm{g}$(MNP)/L) were first diluted with $8\,\mathrm{mL}$ of chloroform and then added dropwise into the polymer solution during sonication for $10\,\mathrm{min}$ to homogenize the mixture. To ensure the homogeneous and efficient intercalation of PMAO-azide polymer hydrocarbon chain with oleic acid on NPs, the mixture was stirred and incubated at room temperature overnight. The ratio of polymer to MNPs corresponds to $500\,\mathrm{polymer}$ units per $nm^2$ of a particle. Next, chloroform was removed very slowly within 5 -6 h through stepwise reduction of pressure to a final value of $350\,\mathrm{mbar}$ and increase of temperature to 34 °C. After a complete removal of chloroform, particles were resuspended in $20\,\mathrm{mL}$ of sodium borate buffer (pH $8.7$) by sonication for $1\,\mathrm{h}$ at 45 °C. The particle suspension was then concentrated to $2\,\mathrm{mL}$ using spin filtration (Amicon regenerated cellulose spin filter, $15\,\mathrm{mL}$, $30\,\mathrm{kDa}$ cutoff size) at $3200\,\mathrm{rpm}$ and 20 °C for $20\,\mathrm{min}$.

#### Particle purification and fractionation

To remove polymeric micelles and partile clusters that are formed during the polymer coating, we fractioned MNCs on non-continuous sucrose gradient ($10\,\%$:$40\,\%$:$60\,\%$, from top to bottom, each fraction $4\,\mathrm{mL}$) in centrifuge falcon tube. Typically, 500 µL of polymer-coated particle suspensions in borate buffer were loaded into the sucrose centrifuge tube and centrifuged for $1.5\,\mathrm{h}$ at $4500\,\mathrm{rpm}$ at 4 °C. Next, to collect monomeric MNCs, meaning nanoparticles that are singly coated within a polymeric shell, nanoparticles between $10\,\%$ and $40\,\%$ band, including $10\,\%$ band, were collected using a long needle. The sucrose was then removed by $3\,\mathrm{rounds}$ of Amicon filtration (Amicon spin filter, $15\,\mathrm{mL}$, $30\,\mathrm{kDa}$ cutoff size) at $3000\,\mathrm{rpm}$ and $20\,^{\circ}\mathrm{C}$ for $15\,\mathrm{min}$. After each round of spin filtration the particles were thoroughly resuspended in borate buffer by vigorous pipetting. Finally, the particles were concentrated to $2\,\mathrm{mL}$ in TE ($5\,\mathrm{mmol}$ Tris, $1\,\mathrm{mmol}$ EDTA, $5\,\mathrm{mmol}$ NaCl, pH 7.3) buffer by spin filtration. To further remove any nonmagnetic impurities from the particles, we performed one round of magnetic washing using a $1.5\,\mathrm{mL}$ MACS column (Miltenyi Biotec) in combination with a MiniMACS separator (permanent magnet) as previously described [2].

#### Agarose gel electrophoresis to purify polymer coated NPs

To extract the fraction of monomeric MNCs from the mixture of polymer coated particles, native agarose gel electrophoresis (AGE) was applied. In a typical preparation procedure, $1.5\,\%$ agarose gel in $1\,\times$ TAE running buffer was made by dissolving $0.5\,\mathrm{g}$ of agarose in 50 mL of $1\,\times$ TAE buffer by $2\,\mathrm{min}$ microwave irradiation. After cooling down at room temperature for $5\,\mathrm{min}$, the homogenous gel solution was casted on a gel mold and left to solidify for $30\,\mathrm{min}$. Afterwards, the casted gel was placed in electrophoresis chamber (Biorad) and samples were pipetted (containing $20\,\%$ of $40\,\%$ sugar solution) into pockets. Next, AGE was run at $90\,\mathrm{mA}$ for $35\,\mathrm{min}$ to separate the monomeric MNCs from the multimeric NPs. Finally, the front section (3-4 mm wide) of the running band that contains monomeric single coated MNCs was excised from the gel. The single particles were then collected by squeezing the gel fragment.

#### DNA labeling of MNCs

Via click-chemistry conjugation, we are able to label DNA single strands onto our MNCs. Here, we labelled our MNCs with 20 nt oligos(dT) that are complementary to 8 nt oligos(dA) on the binding sites of DNA origami. In a typical process, a 150 µL of MNC suspension in TE buffer at a particle concentration of 15 nM was mixed with 32.8 µl of 100 µM 5'-T(20)-DBCO-3' ssDNA in TE to obtain the nominal grafting density of 0.12 ssDNA/nm$^2$. After mixing, the sample was incubated at room temperature overnight. To maximize the binding efficiency of ssDNA to MNCs, we applied a so-called salt aging procedure using 5 M NaCl salt the next day. We increased the concentration of NaCl in the mixture incrementally by 100 mM every hour up to 400 mM. After each salt adjustment step, the mixture was homogenized by vortexing and sonicating for 20 s. At the end of the salt aging procedure, the mixture was again incubated at room temperature overnight. On the next day, the excess of ssDNA was washed out by 2 rounds of centrifugation at 13000 rcf at 5 °C for 12 min. Finally, the sample volume was adjusted back to 150 µl to obtain the original particle concentration. The samples were stored at 4 °C prior to further use.

### 2. STRUCTURAL AND MAGNETIC CHARACTERIZATION OF COLLOIDAL MNCS

#### Dynamic light scattering (DLS)

DLS measurements were performed using a Malvern Zetasizer instrument at 173° backscattered measurement mode. Measurements were performed on 70 µL of particle suspensions in TE buffer (pH 7.3) at a typical particle concentration of 0.015 - 0.02 g/L at 295 K.

#### Magnetic hysteresis loops: characterization of saturation magnetization, magnetic coercivity, and magnetic anisotropy

The magnetic hysteresis loops were measured using magnetic property measurement system (MPMS, Quantum Design). The samples were prepared by pipetting a 100 µL suspension of CMPs in a 6 cm NMR tube. The hysteresis loops were recorded at 298 K using DC scan mode over magnetic fields between -7 and 7 T. The applied magnetic fields were corrected for a remanence field remaining in the chamber and the magnets by referencing measurements against a palladium standard sample. The coercive fields were corrected accordingly.

#### Determination of effective magnetic anisotropy constant

The effective anisotropy constant $K_{\mathrm{eff}}$ is determined using the modified Stoner-Wohlfarth (SW) model proposed by Garcia-Otero et al.[6] The model considers MNPs with uniaxial magnetic anisotropy, randomly distributed magnetization easy axes, and thermal fluctuations. In the classical SW model for randomly distributed MNPs, the coercive field holds $H_{\mathrm{c}}$ = 0.479 $H_{\mathrm{a}}$, in which $H_{\mathrm{a}} = 2K_{\mathrm{eff}}/(\mu_0 M_{\mathrm{s}})$ is the anisotropy field. By taking effects of temperature into account via the Néel relaxation, the temperature-dependent $H_c$ (T) is given by

$$\mu_0 H_c(T) = \mu_0 H_a(T)\left(0.479 - 0.81\left(\frac{k_B T}{2K_{eff}(T)V}\left(\ln\frac{\tau_m}{\tau_0}\right)\right)^{\frac{3}{4}}\right) \quad (1)$$

where $K_{\text{eff}}$ is the effective anisotropy constant and $\mu_0 H_c$ is linearly proportional to $T^{\frac{3}{4}}$ up to certain temperature (Fig. S1a). Assuming $\tau_0 = 10^{-9}$ s and $\tau_m$=100 s, $K_{\text{eff}}$ was computed and plotted over temperature by fitting $\mu_0 H_c$ versus $T^{\frac{3}{4}}$ with Eq.1 (Fig. S1b).

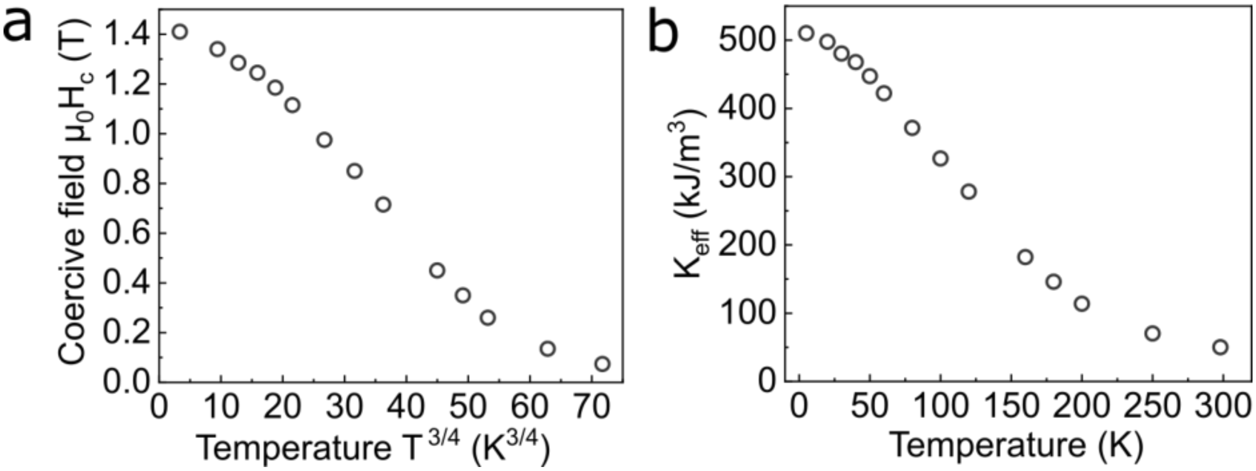


FIG. S1. **Experimental determination of the effective magnetic anisotropy constant**. (a) Coercive field ($H_c$) as a function of temperature measured over a broad range of temperture. (b) $K_{\text{eff}}$ versus temperature.

#### Particle size analysis

Particle size histograms of uncoated (oleic acid coated) and polymer coated MNCs were obtained from the analysis of transmission electron microscopy (TEM) images (JEOL, 100 kV). Negative-stain TEM images were obtained via staining with 1 % uranyl formate, as described in our previous work [2].

### 3. FOLDING AND ANALYSIS OF DNA ORIGAMI STRUCTURES

#### DNA origami folding and purification

6HB DNA origami structures were designed using cadnano 2.5 [7]. The scaffold used for the rotors was the standard $7,249$ bases scaffold, derived from the M13Mp18 bacteriophage. Detailed cadnano designs and sequences can be found in Supplementary Note 1. The 6HBs have a thickness of 6 to 8 nm, and a persistence length of 1.88 to 2.7 µm, showing their robustness and stiffness as mechanical rotation template.

The rotor samples were folded using $100\,\mu\text{L}$ of $100\,\text{nM}$ scaffold (in-house produced in $ddH_2O$) combined with a $5\times$ molar excess of staples (obtained from IDT at $100\,\mu\text{M}$ in 1x IDTE buffer) over the scaffold. Sample buffer was adjusted to 1xTE, $12\,\text{mM}$ $MgCl_2$ and $5\,\text{mM}$ NaCl for folding. Folding was performed by a temperature ramp from $70$ to $40\,°\text{C}$ using a thermal cycling device. The temperature was reduced by $0.1\,°\text{C}$ every $2\,\text{min}$ during the thermal ramp.

Rotor samples were purified in a step-wise manner. Folded samples were at first purified from excess staples using PEG-precipitation similar to reference [8]. Afterwards Neutravidin and fluorescent dyes were added in $10\times$ molar excess over the initial scaffold concentration of the sample during folding. Lastly agarose gel electrophoresis ($70\,\text{V}$,

$1\,\mathrm{hour}$, $0.5\ \times\ TBE$, $5\,\mathrm{mM}$ $MgCl_2$ running buffer) was performed to separate rotor structures from Neutravidine and excess dye, followed by gel extraction to recover the structures from the gel (cf. reference [9]). Extraction from the gel was possible based on the fluorescence from the added Atto655 dyes.

#### Origami structure verification

For the structural verification of the functional assembly between MNPs and DNA origami rotors we used TEM. Negative stain-TEM micrographs were recorded similar to our previous studies (cf. reference [10]), with the addition of a 2-10x excess of MNPS over DNA origami rotor binding sites on the TEM grid. The TEM used for acquisition was a Philips CM-100 at $100\ kV$ and samples were applied to FCF400-CU copper grids from Electron Microscopy Sciences.

#### Fluorescence microscopy sample preparation

The TIRF microscope used in this study is the same as in reference [9]. For measurements the rotor structures were pipetted onto a PEG-modified glass slide (PEG modification protocol cf. reference [11]) at a concentration of $200\,\mathrm{pM}$. The measurement buffer used during the fluorescence acquisistions was made up from $0.3 \times$ TB ($30\,\mathrm{mM}$ Tris, $30\,\mathrm{mM}$ Borate), $3\,\mathrm{mM}$ $MgCl_2$ in $ddH_2O$ with 48 % sucrose. The sample chambers used for fluorescence measurements were similar to previous study and formed a $50\,\mathrm{\mu m}$ high channel of $12\,\mathrm{mm}$ length. After the sample was applied to the PEG-biotin surface, the chamber was washed with at least $5\ \times$ total volumes of the measurement chamber. After the origami were bound to the surface, MNPs were added in excess by flushing and incubation of $10\,\mathrm{\mu L}$ ($10\,\mathrm{nM}$ MNPs in $1 \times$ TE $20\,\mathrm{mM}$ $MgCl_2$) in the measurement chamber for 30 minutes. For the measurement, the chamber and wells were filled with measurement buffer and buffer was refilled regularly to restrict evaporation caused changes in the sample chamber.

#### Homogeneous and rotating magnetic fields

The external magnetic fields we applied during this study, were generated using two pairs of pseudo-Helmholtz coils, which were placed in a cross-wise arrangement around the sample chamber and controlled using a pair of Amplifiers.

#### Analysis of single-molecule fluorescence measurements

Fluoresecence data acquired using our single-molecule TIRF microscopy setup tracking measurements was processed using the *Picasso* software package from reference [12]. In all of the data acquisitions, we used the $42$ fluorescent dyes at the tip of the rotor structure for single-molecule structure tracking. Therefore the combined fluorescence of all dyes was used for the positional localization of the rotor structures during every timestep of the measurement. If movements of the sample or sample stage changed the emitter position during measurements, a drift correction function of the *Picasso* package was applied to realign the datapoints.

In a next step the data was imported for further analysis using a custom python analysis routine. In that routine the center of each rotor was determined using a circular fit of the acquired data. This was achieved using a circular fit. In a next step, the positional coordinates of all localizations were transformed into relative polar coordinates per rotor. Rotor trajectories were generated by plotting the normalized rotor position as a function of average time.

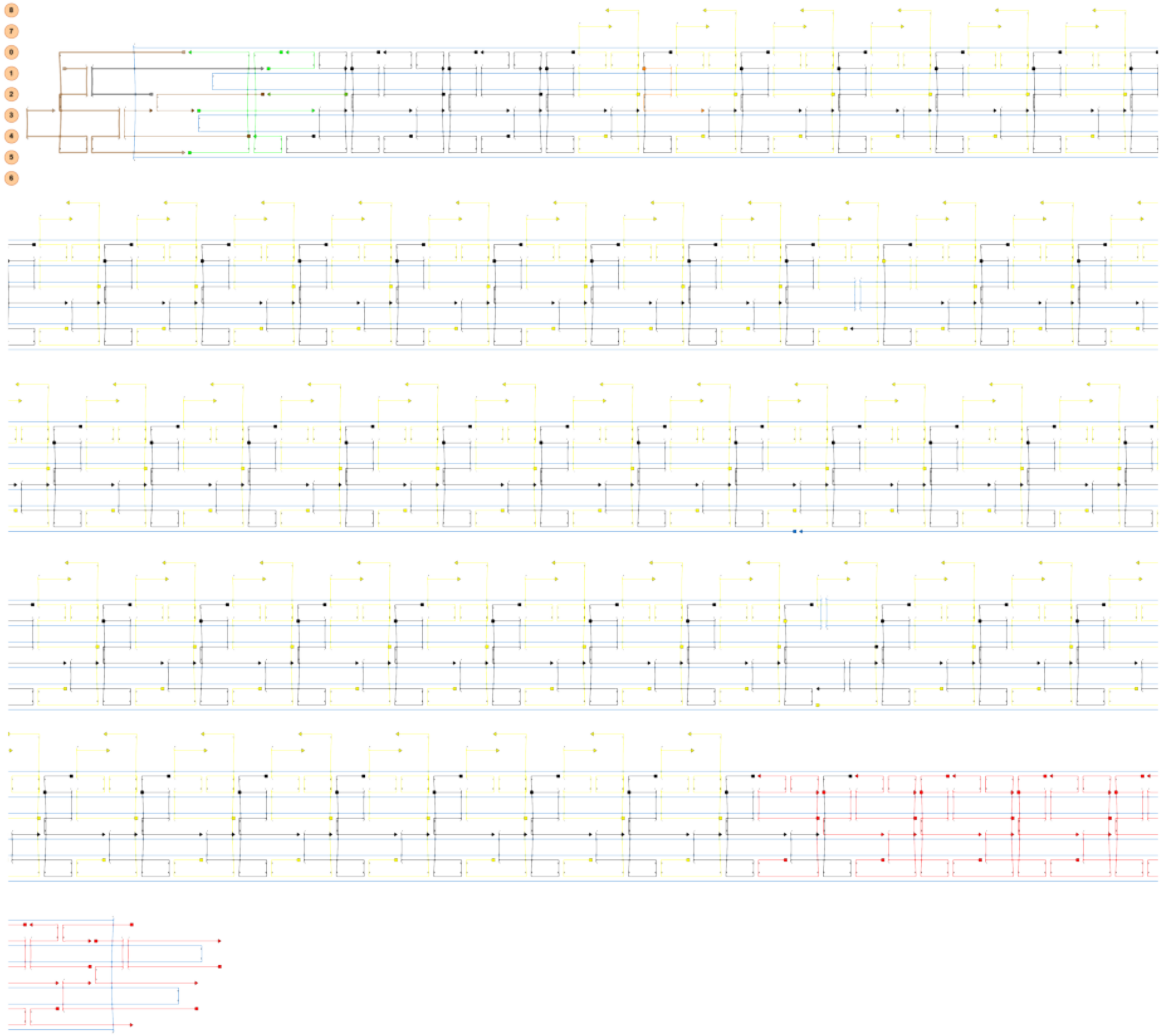

FIG. S2. **DNA origami design of multi-MNC MADONAs**. Staple colors show special functionalities. Black = unmodified basic staples. Red = Staples with extension sequence to bind fluorescent dyes. Brown and green = blunt end design. Yellow = Staples extended on helix 0 and 1 with 8 nt oligos(dA) to bind MNCs. Orange = Biotin modified staple site.

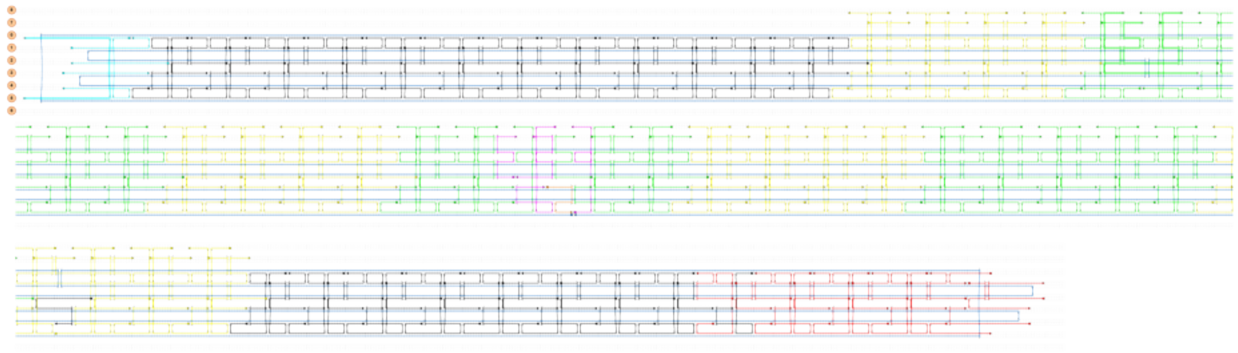

FIG. S3. **DNA origami design of 4-MNC MADONAs**. Staple colors show special functionalities. Black = unmodified basic staples. Red = Staples with extension sequence to bind fluorescent dyes. Cyan = Passivated end design of 6hb. Yellow = Staples extended on helix 0 and 1 with 8-nt Poly-A to bind MNCs for structures with 1 to 4 MNCS. Green = Yellow = Staples extended on helix 0 and 1 with 8 nt oligos(dA) to bind 4 MNCs to the bundles. Magenta= modified staple routing for Biotin mid pivot. Orange = Biotin modified staple site.

### 4. TRAP STIFFNESS DATA FOR MULTI-MNC MADONAS

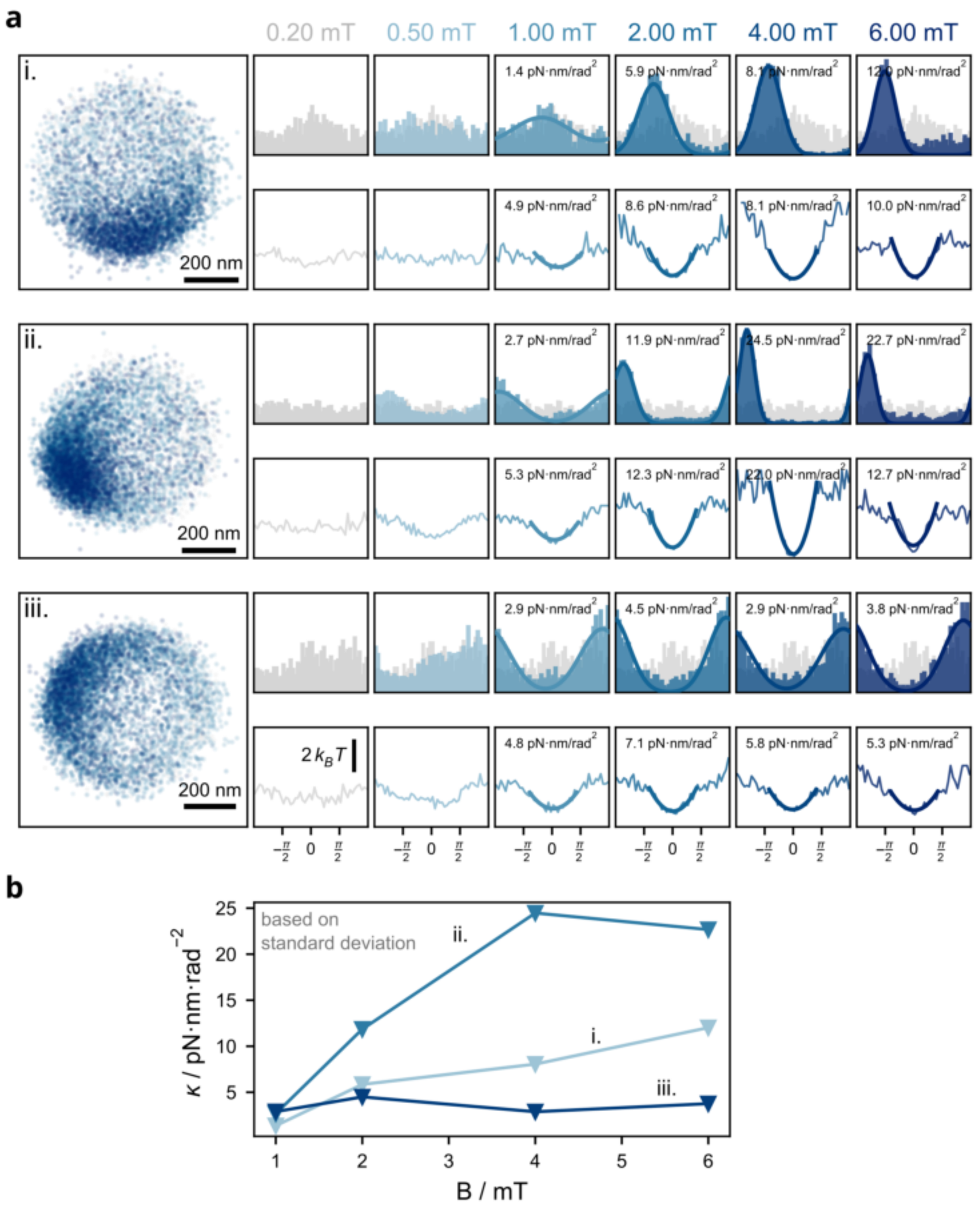


FIG. S4. **Trap stiffness of Multi-MNC MADONAs during clamping experiment**. (a) Localization histograms (top row) and energy landscapes (bottom row) for exemplary particles overlayed with a Gaussian or quadratic fit estimating the trap stiffness. (b) Angular trap stiffness $\kappa$ as a function of field strength $B$ based on the Gaussian fits.

### 5. TEM MICROGRAPHS OF ROTORS DECORATED WITH MAGNETIC NANOPARTICLES

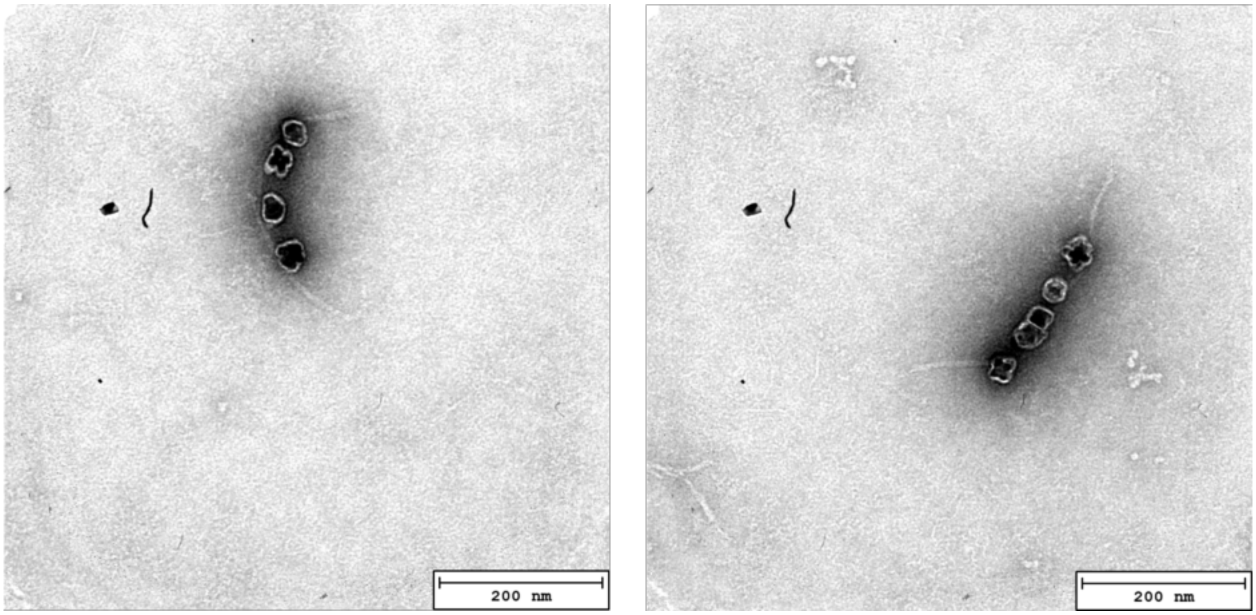


FIG. S5. **Negative-stain TEM micrographs of multi-MNC MADONAs**. These rotors were prepared through coupling of MNCs on the 6HBs in solution. The structures were then purified from unbound MNCs and 6HBs using excise-squeeze technique. In the multi-MNC design, the particle binding sites are placed 6-8 nm apart from each other, binding of several particles with the center-to-center distance of $\approx$ 17 nm are expected. This means the maximum number of MNCs per rotors is seven to eight. However, these TEM images show that, persumably due to the fact that the binding sites are very closed to each other, the actual number of bound MNCs per structure is far from ideal.

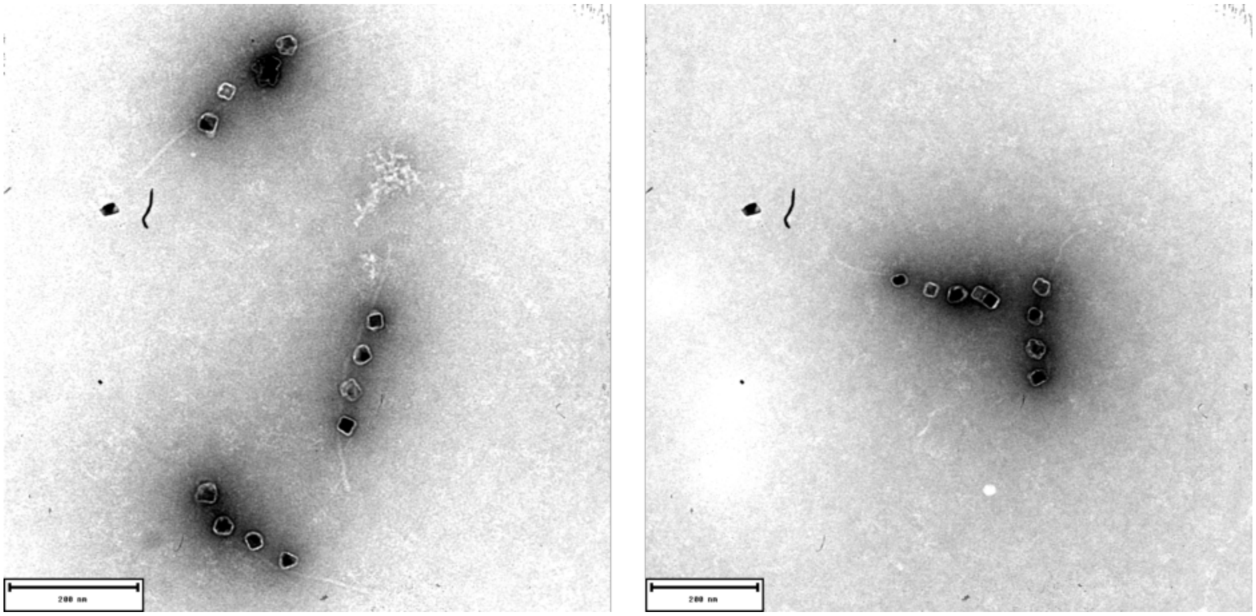


FIG. S6. **Negative-stain TEM micrographs of 4-MNC MADONAs**. These rotors were prepared by coupling of MNCs on the 6HBs in-solution. The structures were then purified from unbound MNCs and 6HBs using excise-squeeze technique. In the 4-MNC design, the particle binding sites are placed at 64 nm ($R_{\mathrm{c-c}}$) distance. TEM images of five exemplary nanorotrs show a good quantitative binding of 4 MNCs to 4 binding sites for each structure.

### 6. CO-ROTATION AS PHASE-LOCKING PHENOMENA

We consider a DNA origami rotor with magnetic nanocubes rigidly attached to its structure, while each magnetic cube carries a magnetic dipole moment $m_i$. Then, the resulting magnetic moment of the rotor $m_{\text{eff}}$ experiences a torque $\tau_m = m_{\text{eff}} B \sin(\varphi_B - \varphi_m)$, depending on its axis' orientation $\varphi_m$ relative to the external field $\varphi_B$. Since the magnetic particles are rigidly coupled to the DNA origami, the rotor co-rotates, and there is a constant angle offset between $m_{\text{eff}}$ and the rotor axis, $\varphi_m = \varphi_r + \Delta\varphi$. For notational simplicity, we eliminate the constant phase offset by setting $\Delta\varphi = 0$, thereby virtually aligning the rotor axis with the axis of the magnetic moment. The rotor trajectory for a rotating magnetic field at constant angular velocity, $\varphi_B(t) = \omega t$, is described by a torque balance in the overdamped regime, where the induced magnetic torque has to compensate for the viscous drag, the local restoring torque stemming from rotor-surface interactions, and thermal fluctuations.

$$\zeta \dot{\varphi}_r + U'(\varphi_r) + \xi(t) = m_{\text{eff}} B \sin\big(\omega t - \varphi_r\big). \tag{2}$$

Here, $\zeta$ is the drag coefficient, $U(\varphi_r)$ the energy landscape, and $\xi$ Gaussian white noise with $\langle \xi(t)\,\xi(t')\rangle = 2\zeta k_B T\,\delta(t-t')$. We switch to a rotating-frame description by defining a phase offset between the magnetic field and the rotor's angular position,

$$\theta = \omega t - \varphi_r, \quad \dot{\theta} = \omega - \dot{\varphi}_r. \tag{3}$$

Then, substituting $\varphi_r$ into torque balance yields

$$\dot{\theta} = \omega - \frac{m_{\text{eff}}\, B}{\zeta} \sin\theta + \frac{1}{\zeta} U'\big(\omega t - \theta\big) + \sqrt{2D}\,\eta(t), \tag{4}$$

with a normalized noise term $\frac{1}{\zeta}\,\xi(t) = \sqrt{2D}\,\eta(t)$, $D$ being the rotational diffusion coefficient. We further assume the energy landscape of the rotor to be $U(\varphi_r) = -\Delta E \cos(n\varphi_r)$, which leads to an expression for the surface-induced torque of $U'(\omega t - \theta) = n\Delta E \sin\big(n(\omega t - \theta)\big)$ and we arrive at

$$\dot{\theta} = \omega - \frac{m_{\text{eff}}\, B}{\zeta} \sin\theta + \frac{n\Delta E}{\zeta} \sin\big(n(\omega t - \theta)\big) + \sqrt{2D}\,\eta(t). \tag{5}$$

Finally, denoting $K = m_{\text{eff}}\, B/\zeta$ as the coupling strength and $\Delta E' = \Delta E/\zeta$ as the scaled surface-interaction potential, the phase obeys a generalized stochastic Adler equation

$$\dot{\theta} = \omega - K\,\sin\theta + n\Delta E'\,\sin\big(n(\omega t - \theta)\big) + \sqrt{2D}\,\eta(t), \tag{6}$$

where the positive $\Delta E'$-term represents the additional desynchronizing effect of the local surface interactions. Altogether, Eqn. (6) describes a non-autonomous stochastic Adler–type system – mathematically equivalent to an overdamped Brownian particle in a time-varying washboard potential. The system reveals regimes of sustained phase locking for strong coupling, intermittent phase slips, and thermally activated escapes at lower coupling strength relative to the detuning term $\omega$.

**Classical Adler equation**

In the limit of a flat energy landscape ($\Delta E' \to 0$) and vanishing fluctuations ($\eta = 0$), Eqn. (6) reduces to the classic Adler equation,

$$\dot{\theta} = \omega - K\,\sin\theta, \tag{7}$$

which describes the deterministic synchronization (of a natural oscillator) to an external periodic drive. Since our system has no intrinsic frequency, the detuning term is simply $\omega$. The rotors are phase-locked to the external drive, $\dot{\theta} = 0$, if the magnetic torque compensates the viscous drag. In the classic Adler equation, a fixed point exists whenever

$$|\omega| \leq K \quad \text{and} \quad m_{\text{eff}}\, B \geq \zeta\,|\omega|, \tag{8}$$

which implies that the rotors' magnetic moment must satisfy $m_{\text{eff}} \geq \zeta|\omega|/B$. The constant phase difference in the locked state is then given by $\arcsin(\omega/K)$. Equivalently, one may view $\theta(t)$ as the position of a particle in a washboard potential

$$V(\theta, t) = -\,\omega\,\theta - \,K\cos\theta, \tag{9}$$

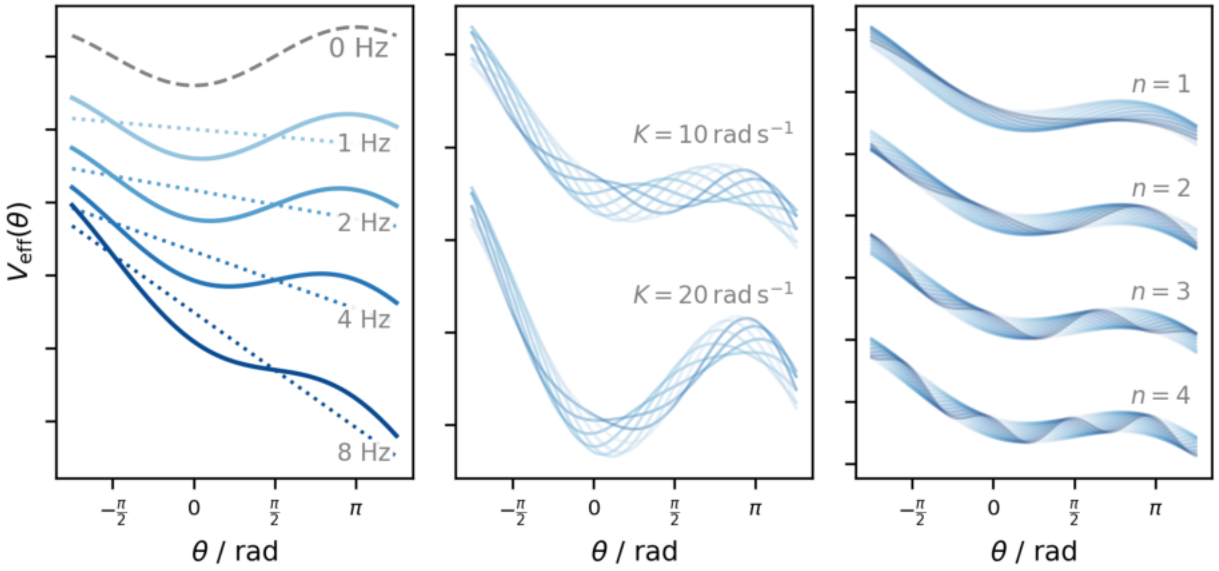


FIG. S7. **Effective potential $V_{\rm eff}(\theta)$ and locking regimes under different conditions**. (a) Classic Adler potential in case of $\Delta E' = 0$: $V_{\rm eff}(\theta) = -\omega\,\theta - K\cos\theta$ traps the phase in its cosine wells at low drive frequency $f$, whereas at high $f$ the linear tilt $-\omega\theta$ dominates and phase slipping occurs. (b) The effect of a periodically explored surface-interaction $U(\varphi_r)$ leads to a time-dependent corrugation of $V_{\rm eff}(\theta)$. Depending on the surface interaction strength, the coupling $K$ might not be sufficient to keep the rotor phase-locked ($\Delta E = 4\,k_BT$, $n = 2$, $f = 1$ Hz). The weak coupling yields a shallow well that slips intermittently under the tilt. (c) The number of wells $n$ in the surface interaction potential $U(\varphi_r)$ affects the corrugation of $V_{\rm eff}(\theta)$ and allows fractional phase locking by providing additional stable wells ($\Delta E = 4\,k_BT$, $f = 1$ Hz, $K = 10\,{\rm rad\,s^{-1}}$).

as illustrated in Fig.S7a. In case of sufficient coupling, the particle remains trapped in a local minimum despite the overall tilt – this corresponds to a constant phase difference and the regime of phase locking. However, if $|\omega| > K$, the tilt overwhelms the wells and the particle runs downhill with an average drift rate

$$\left\langle \dot{\theta} \right\rangle \;=\; \mathrm{sign}(\omega)\sqrt{\omega^2 - K^2}. \tag{10}$$

In the lab frame this means that the rotor can no longer keep up with the applied field and instead *slips* in phase, see Fig. S10.

**Stochastic Adler equation**

If we now allow noise to kick in, $D > 0$ but $\Delta E' = 0$, we observe features of a noisy synchronizer. Then, the Adler equation reads $\dot{\theta} = \omega - K\sin\theta + \sqrt{2D}\,\eta(t)$, where phase locking can be thought of as a diffusing virtual particle $\theta$ trapped in a potential well that represents the locked state, see Eqn. 9. However, random thermal kicks occasionally give the particle enough energy to hop over the barrier $\Delta V$ into the next well. In the Kramers picture, escape events scale with

$$k \;\propto\; \exp\!\left(-\frac{\Delta V}{D}\right). \tag{11}$$

If $D$ is small, the escapes are rare, and we can observe long periods of phase-locking under conditions deterministic synchronization would occur. Exemplary trajectories are shown in Fig. S9

**Breathing potential**

If we additionally consider the orientation-dependent local interactions of the DNA origami rotor with the surface, $\theta(t)$ experiences an periodic forcing term $n\Delta E'\,\sin\!\big(n(\omega t - \theta)\big)$, which leads to a 'breathing' energy landscape in the rotating frame

$$V(\theta, t) = -\,\omega\,\theta -\; K\cos\theta -\; \Delta E'\,\cos\!\big(n(\,\omega t - \theta)\big)\,, \tag{12}$$

where the washboard potential is continuously modulated by the oscillatory component $\Delta E'$, see also Fig. S7b and c. Then $\theta(t)$ cannot settle into a static fixed point anymore, but lives on a periodic orbit

$$\theta(t + qT) \;=\; \theta(t) + 2\pi\, m, \tag{13}$$

with integer number $m$ that closes only after $q$ cycles of the drive, while in the lab frame the rotor angle advances by $\Delta\varphi_r = 2\pi\,(q - m)$. The rotation number

$$N_r \;=\; \frac{\Delta\varphi_r}{2\pi} \;=\; \frac{q - m}{q} \tag{14}$$

then describes the number of full rotor turns per drive cycle.

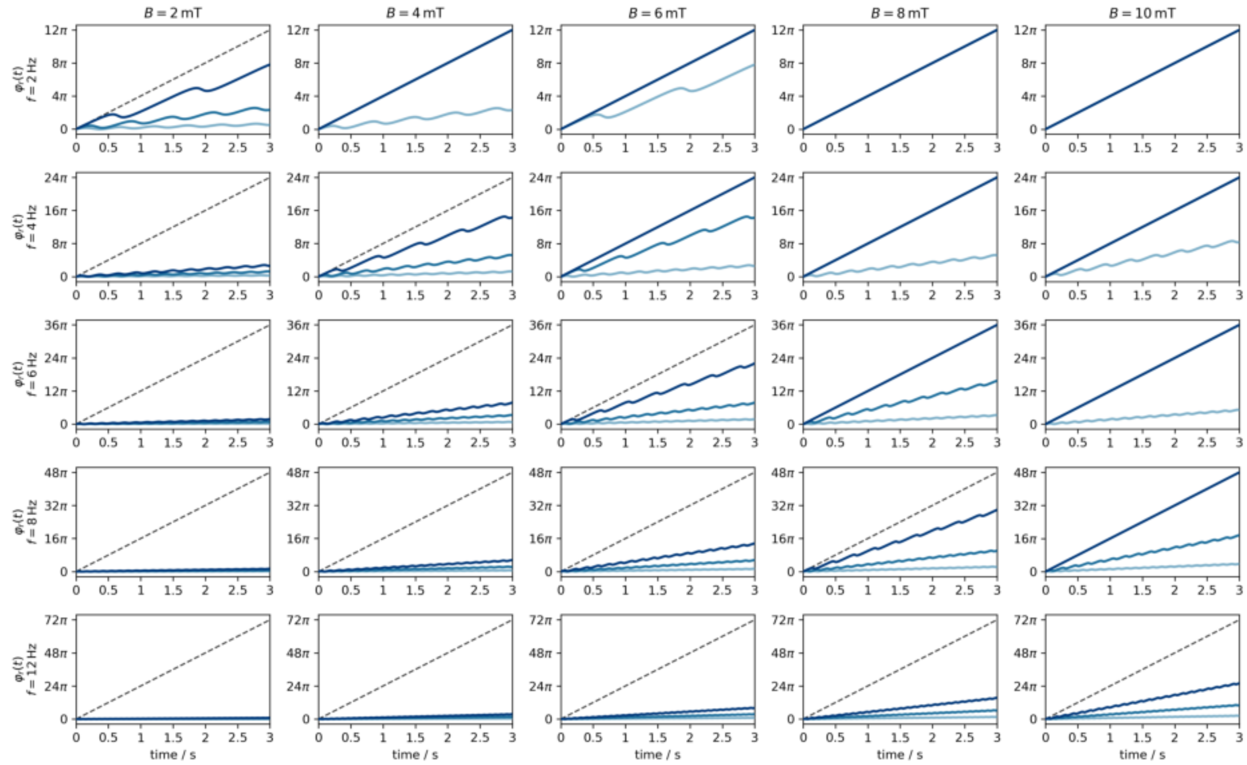


FIG. S8. Deterministic rotor trajectories at different actuation frequencies and field strengths for $D = 0$ and $\Delta E = 0$. Light to dark blue curves correspond to effective magnetic moments $m_{\mathrm{eff}} = \{1, 2, 3\} \cdot 10^{-18}\mathrm{A\,m}^2$.

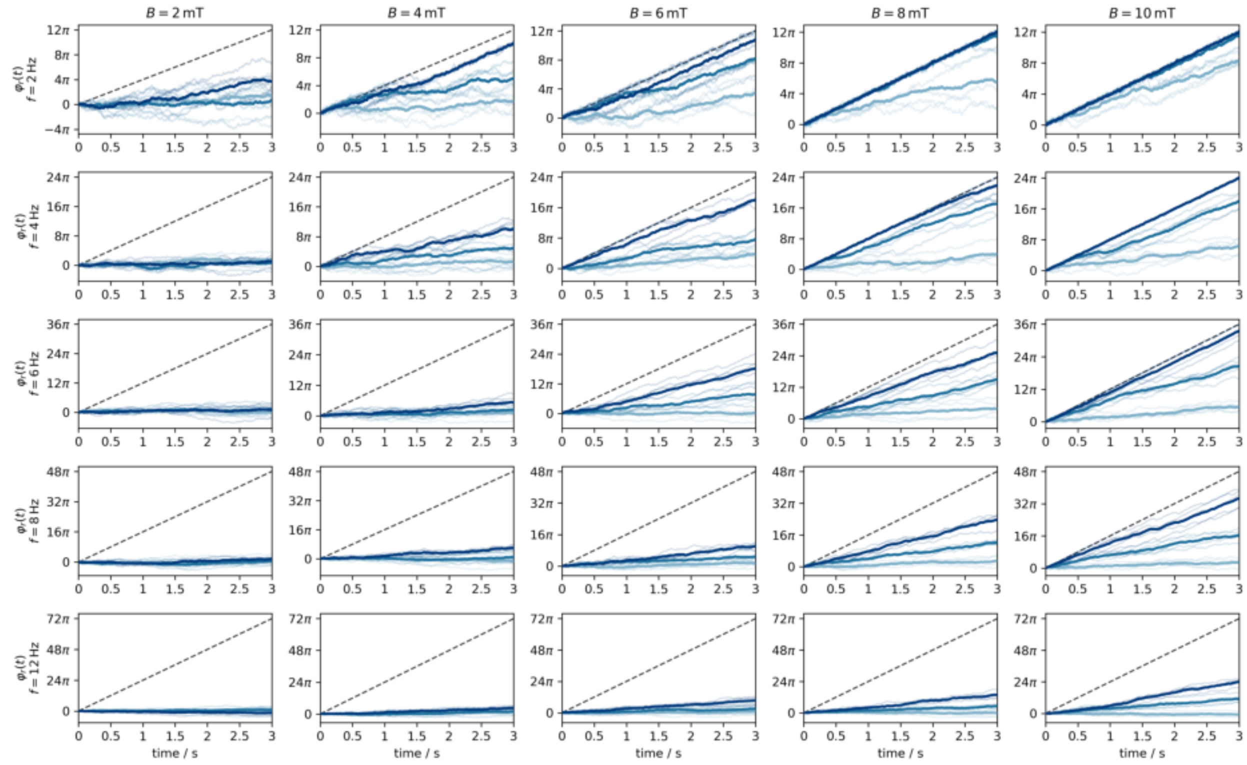


FIG. S9. Stochastic rotor trajectories at different actuation frequencies and field strengths for rotors with a rotational diffusion coefficient $D = 8\,\mathrm{rad}^2\,\mathrm{s}^{-1}$. Light to dark blue curves correspond to effective magnetic moments $m_{\mathrm{eff}} = \{1, 2, 3\} \cdot 10^{-18}\mathrm{A\,m}^2$. For each magnetic moment 5 exemplary trajectories and their mean trend are shown.

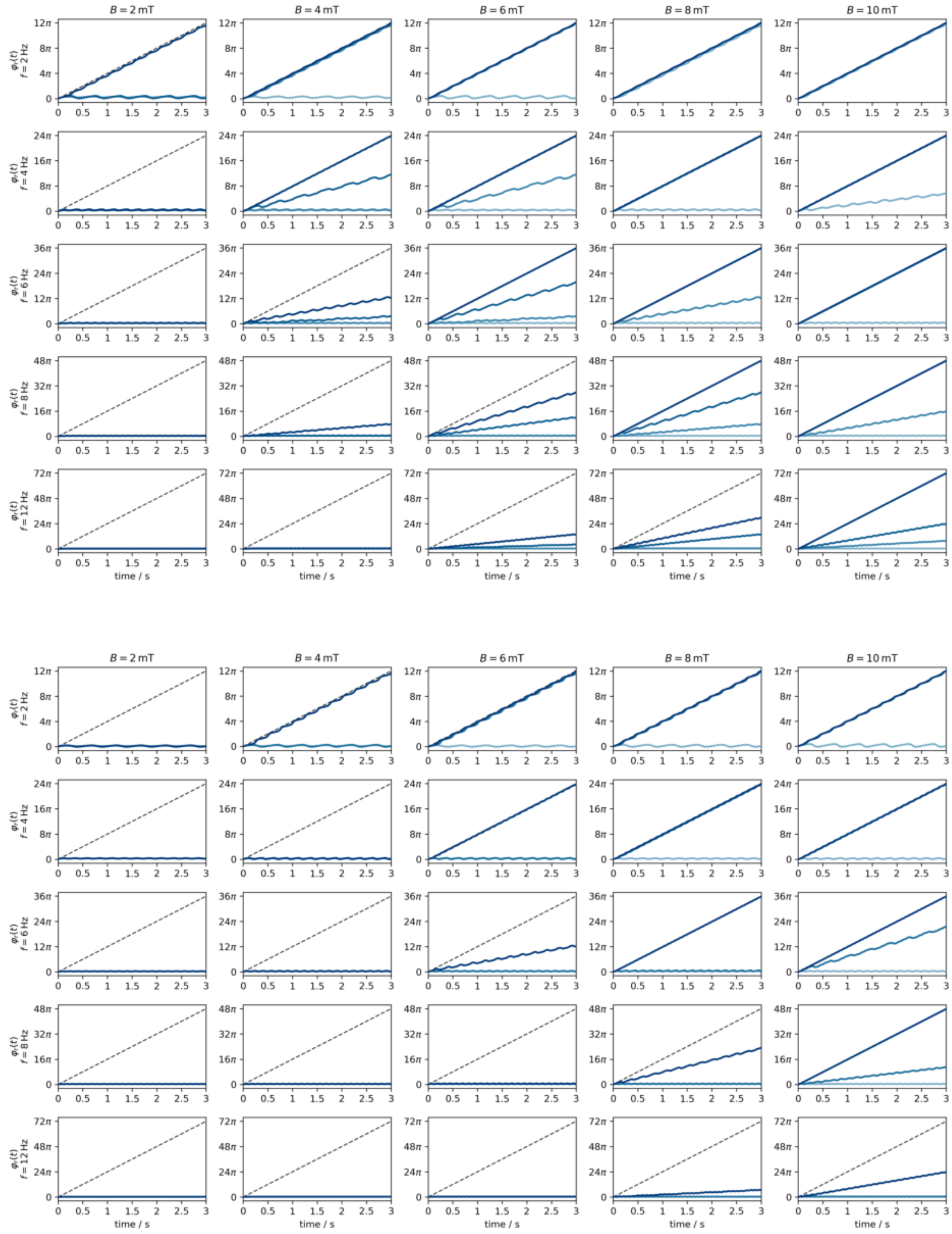


FIG. S10. Deterministic rotor trajectories at different actuation frequencies and field strengths for rotors with an intrinsic sinusoidal $\pi$-periodic energy landscape with $\Delta E = 0.5 k_B T$ (top) and $1 k_B T$ (bottom). Light to dark blue curves correspond to effective magnetic moments $m_{\text{eff}} = \{1, 2, 3\} \cdot 10^{-18}\,\text{A}\,\text{m}^2$.

### 7. DROP-OUT ANALYSIS FOR ROTATIONS AT DIFFERENT ANGULAR VELOCITIES AND FIELD STRENGTHS

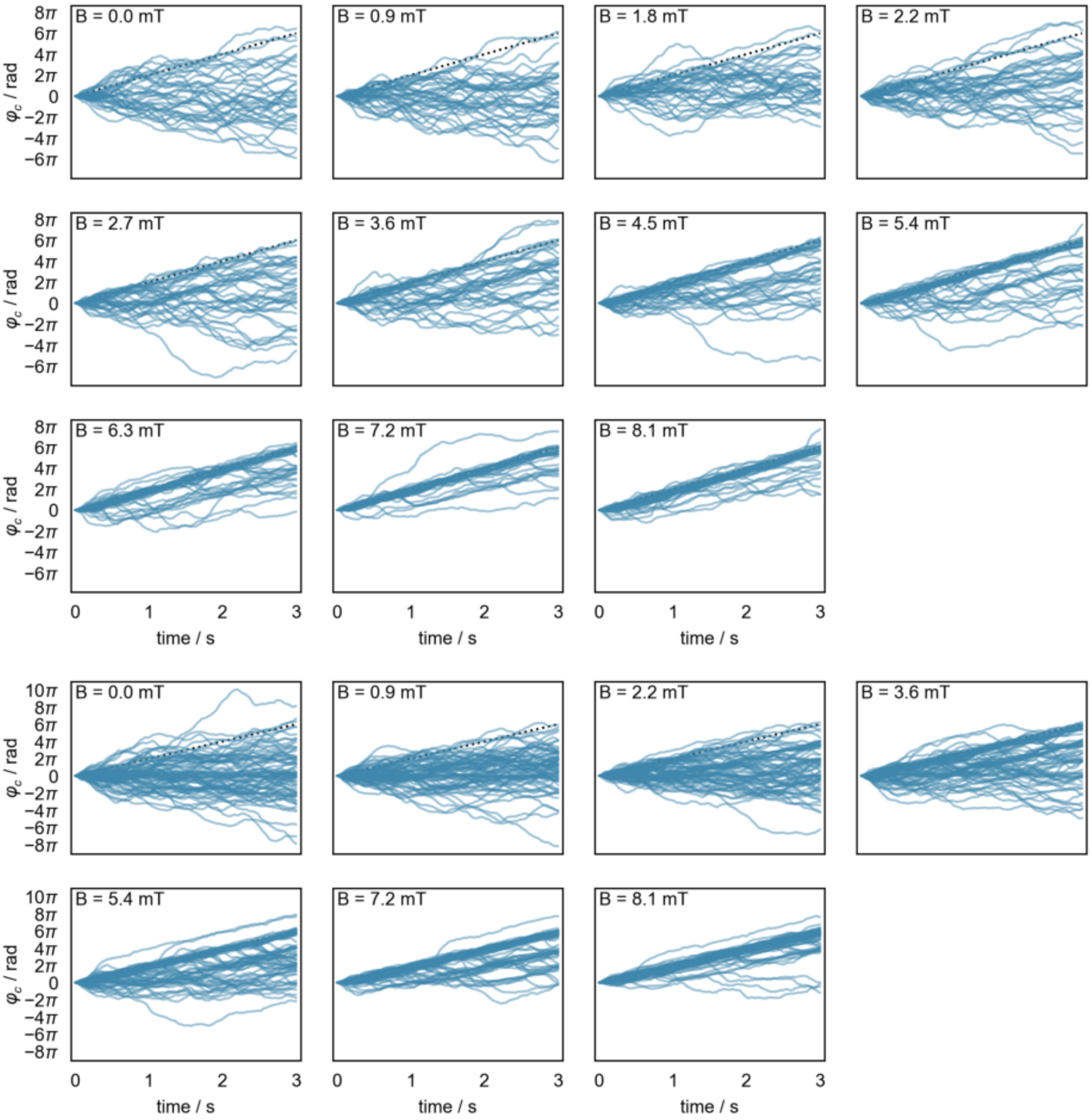


FIG. S11. TIRFM-RMF assay at $1\,\mathrm{Hz}$ corresponding to data shown in Fig. S15c to Fig. S15d. Smoothed rotor trajectories for rotations at $1\,\mathrm{Hz}$ with different field strengths for two different set of rotors. The dashed line corresponds to a perfect synchronization with the field.

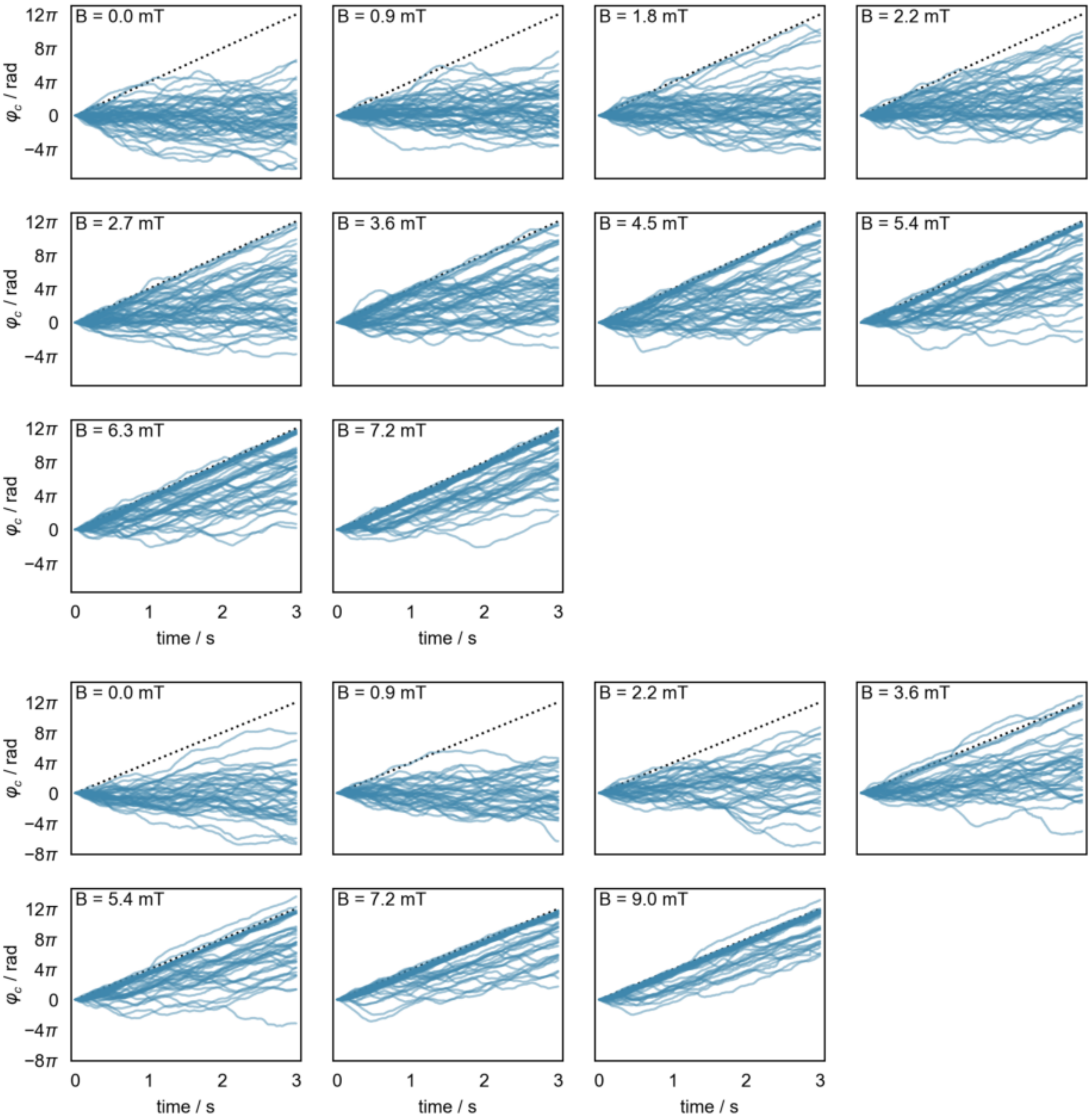


FIG. S12. TIRFM-RMF assay at $2\,\mathrm{Hz}$ corresponding to data shown in Fig. S15c to Fig. S15d. Smoothed rotor trajectories for rotations at $2\,\mathrm{Hz}$ with different field strengths for two different set of rotors.

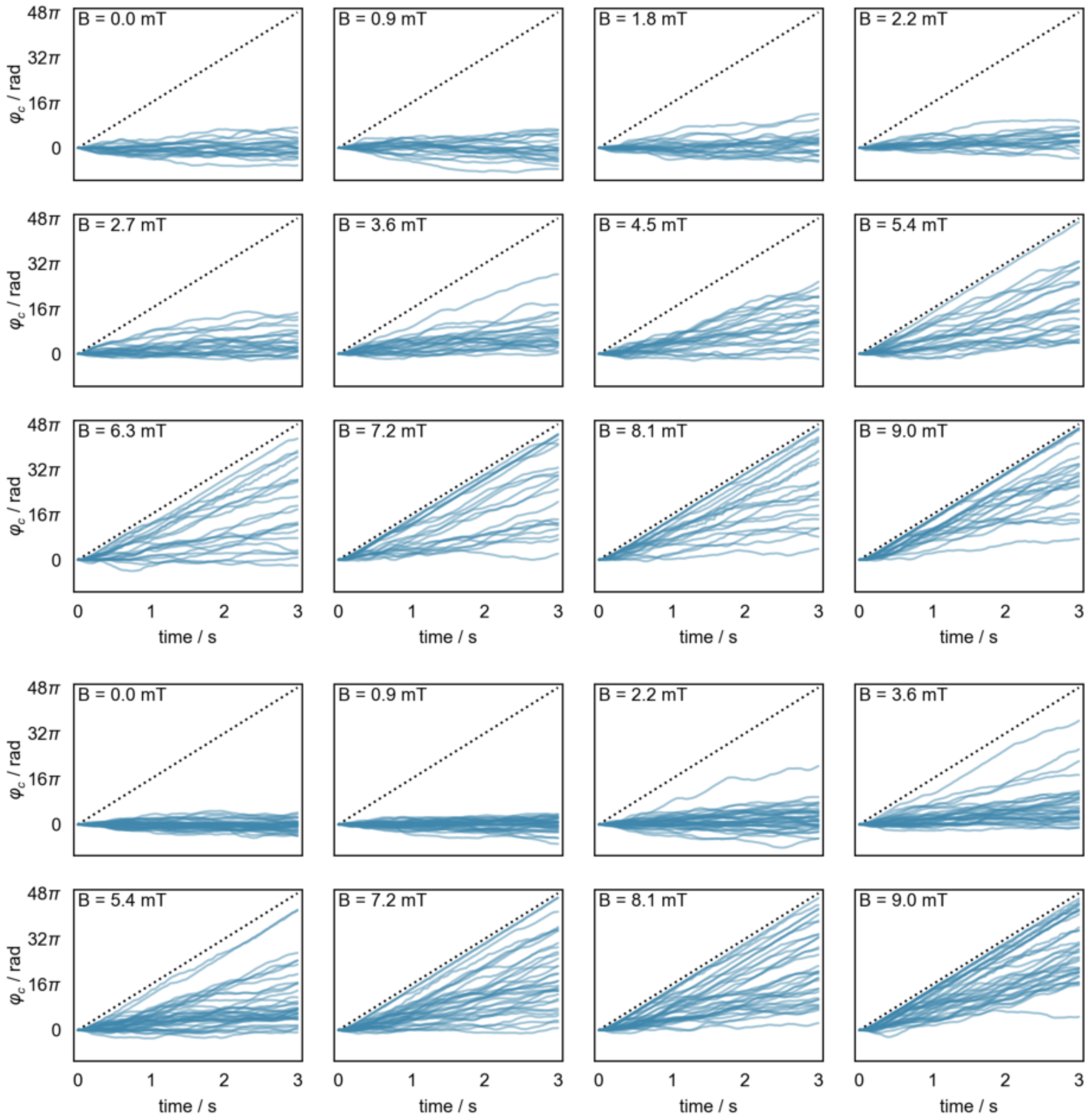


FIG. S13. TIRFM-RMF assay at $8\,\mathrm{Hz}$ corresponding to data shown in Fig. S15c to Fig. S15d. Smoothed rotor trajectories for rotations at $8\,\mathrm{Hz}$ with different field strengths for two different set of rotors.

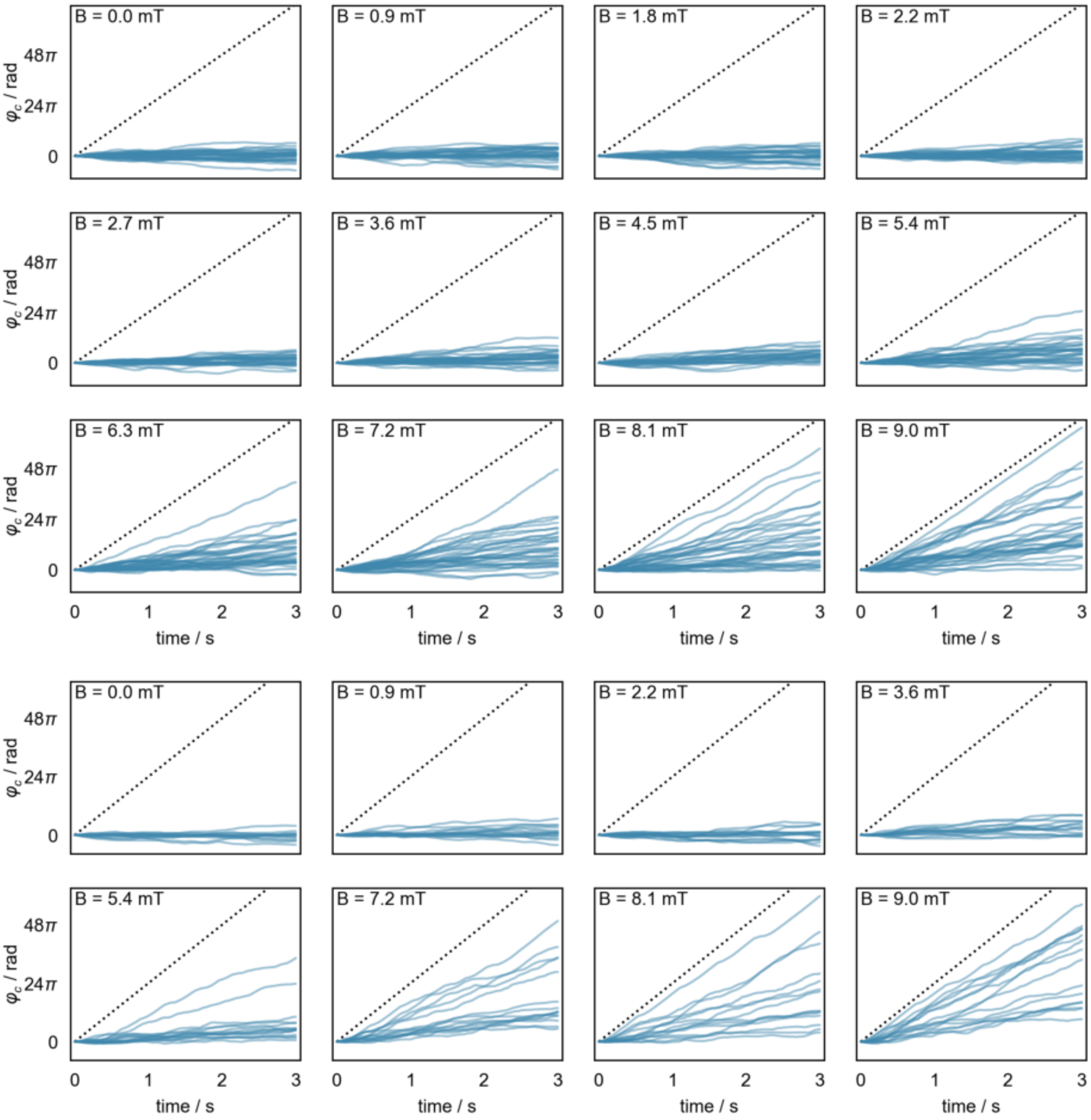


FIG. S14. TIRFM-RMF assay at $12\,\mathrm{Hz}$ corresponding to data shown in Fig. S15c to Fig. S15d. Smoothed rotor trajectories for rotations at $12\,\mathrm{Hz}$ with different field strengths for two different set of rotors.

### 8. MAGNETIC TORQUES OF 4-MNC MADONAS FROM FREQUENCY-RESOLVED DROP-OUT ANALYSES

#### Phase-locking analysis at different frequencies

To quantify the magnetic torques of 4-MNC MADONAs from our rotation assays (Figure 3e and 3g), we computed the so-called phase-locking value (PLV) of 4-MNC MADONAs with respect to the external field given by

$$PLV = \frac{1}{N}\sum_{n=1}^{N} e^{i(\omega_B t_n - \varphi_{r,n})} \tag{15}$$

where PLV = 1 indicates perfect co-rotation with the external field. For instance, Figure 3f shows the temporal evolution and the distribution of PLV for the highlighted rotor (orange-brown) in Figure 3e during actuations at 2 Hz estimated from overlapping 200 ms windows. First, we see that at low field strength, thermal fluctuations significantly affect the rotors' motion and thus the PLV value for these measurements but generally we expect continuous co-rotation to be characterized by PLV > 0.7. However, the PLV values are not directly comparable across different actuation frequencies when the total measurement duration is kept fixed. The reason is that the effective signal-to-noise ratio of phase-locking detection increases with actuation frequency (Figure 3e and Figure 3g), thus for the same recording time and constant $D$, higher-frequency actuation provides more cycles, so coherent phase-locking can be detected more reliably. We therefore performed TIRFM–RMF assays for a given field of view at a fixed actuation frequency (Figure 3e for 2 Hz; Figure 3g for 8 Hz) while varying the magnetic field strength to determine the breakdown points of the rotor ensemble. For both actuation conditions, the fraction of phase-locked rotors increases markedly with the magnetic field strength, which is consistent with stronger magnetic torque that stabilizes the phase locking (Figure S15a and Figure S15b). In addition to the classic torque balance between viscous and magnetic torque, local restoring torques in the rotors' individual energy landscapes can critically interfere and temporally trap the rotor[9, 10]. We therefore established a second criterion based on the PLV values, where we consider a rotor at least temporally following the field synchronously if its mean PLV value exceeds the $1.1 \times \mathrm{PLV}_{\mathrm{diff,max}}$ of the ensemble. Using this PLV criterion, we performed drop-out analyses on the datasets shown in (Figure 3e and 3g of the main text) to estimate $m_{\mathrm{eff}}$ of individual rotors based on the survival curves (Figure S15c), showing the number of rotors for different initial rotor sets that satisfy the PLV criterion as function of the normalized field, $\omega_{\mathrm{B}}$/B. Upon decreasing the field strength for a given $\omega_{\mathrm{B}}$, more and more rotors drop out. Using the inequality for each rotor, we can determine a distribution of a lower bound $m_{\mathrm{eff}}$, ranging from values ($1 \times 10^{-18}$ A $\mathrm{m}^2$) associated with a single MNC up to moments one order of magnitude higher (Figure S15d). Consequently, the torque $\tau$ of a 4-MNC MADONAs varies between few pN nm to > 100 pN nm (Fig. S15e).

Looking more closely at the experimental magnetic moment and torque histograms (Fig. S15d and Fig. S15e), we observe a subset of rotors with torques in the order of 50-100 pN nm. We propose that these are the rotors, in which instead of monomeric MNCs, some dimers and trimers (MNCs that were attached perfectly aligned during the polymer coating) were attached to some binding site. In this situation, and as these dimers/trimers are magnetically well aligned even in the absence of external magnetic fields, their contribution to the overall magnetic moment of the rotors is significant. There is further a subset of rotors with torques below 10 pN nm, which are presumably those rotors having less than 4 MNCs or MNCs with antagonistic starting vectors of $m$.

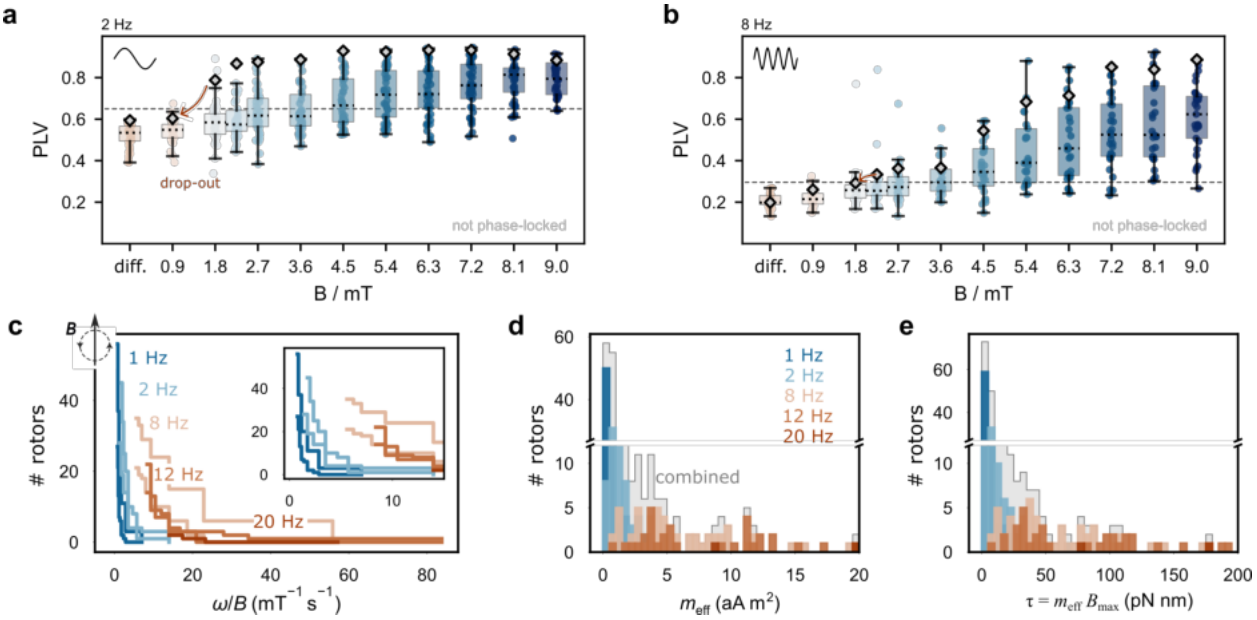


FIG. S15. **Determination of magnetic torques of 4-MNC MADONAs using drop-out analysis and PLV criteria**. (a, b) Mean PLV values of two rotor ensembles across increasing field strengths at 2 and 8 Hz, respectively. Diamond markers denote the orange-highlighted rotors shown in Figure 3e and 3g of the main text. The dashed line marks an actuation threshold defined as 1.1 × $\mathrm{PLV}_{\mathrm{diff,max}}$. We classify a rotor as driven rotor with the RMF if its PLV during actuation exceeds this threshold – e.g., it shows a detectable response above its diffusive baseline—even if it does not closely phase-lock to the rotating field. (c) Population of surviving rotors versus normalized field $\omega$/B ($\mathrm{s}^{-1}$ $\mathrm{mT}^{-1}$). The curves were obtained by tracking the drop-out of rotors at decreasing field strength for giving driving frequencies. The threshold for considering a particle as dropped out was determined relative to the mean PLV value during diffusive recordings. (d) Lower-bound $m_{\mathrm{eff}}$ histograms, where $m_{\mathrm{eff}}$ was inferred from the dropout analysis by using $m_{\mathrm{eff}} = \tau_{\mathrm{drop}} / B_{\mathrm{drop}}$, while $\tau_{\mathrm{drop}} = \zeta_{\mathrm{rot}}\omega$ and $\zeta_{\mathrm{rot}} = k_{\mathrm{B}}T/D_{\mathrm{rot}}$ for each rotor. (e) Magnetic torques $\tau = m_{\mathrm{eff}}\ B_{\mathrm{max}}$ histograms during the experiment based on the effective magnetic moments in (d) and 9 mT.

### 9. TRAP STIFFNESS DATA FOR 4-MNC MADONAS

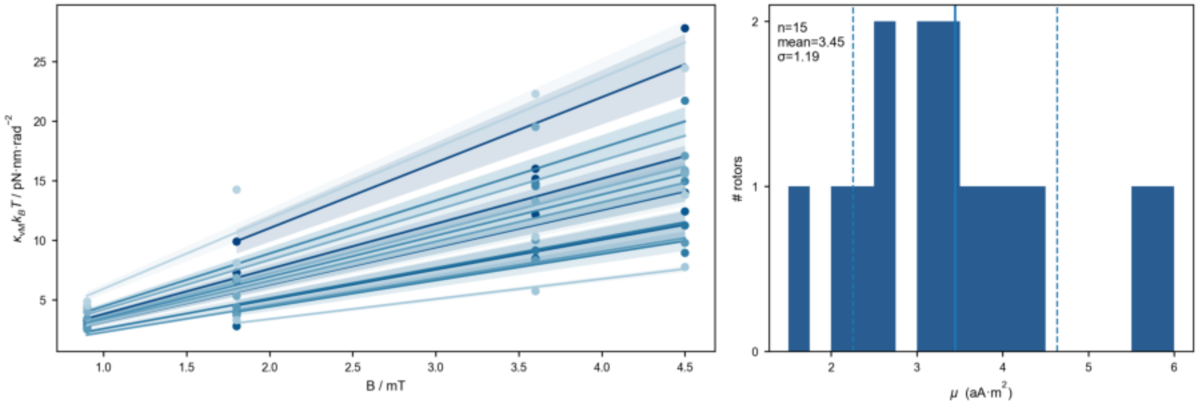


FIG. S16. **4-MNC MADONA clamping experiment**. Traps stiffness as function of the applied field (left) and extracted effective magnetic moment (right).

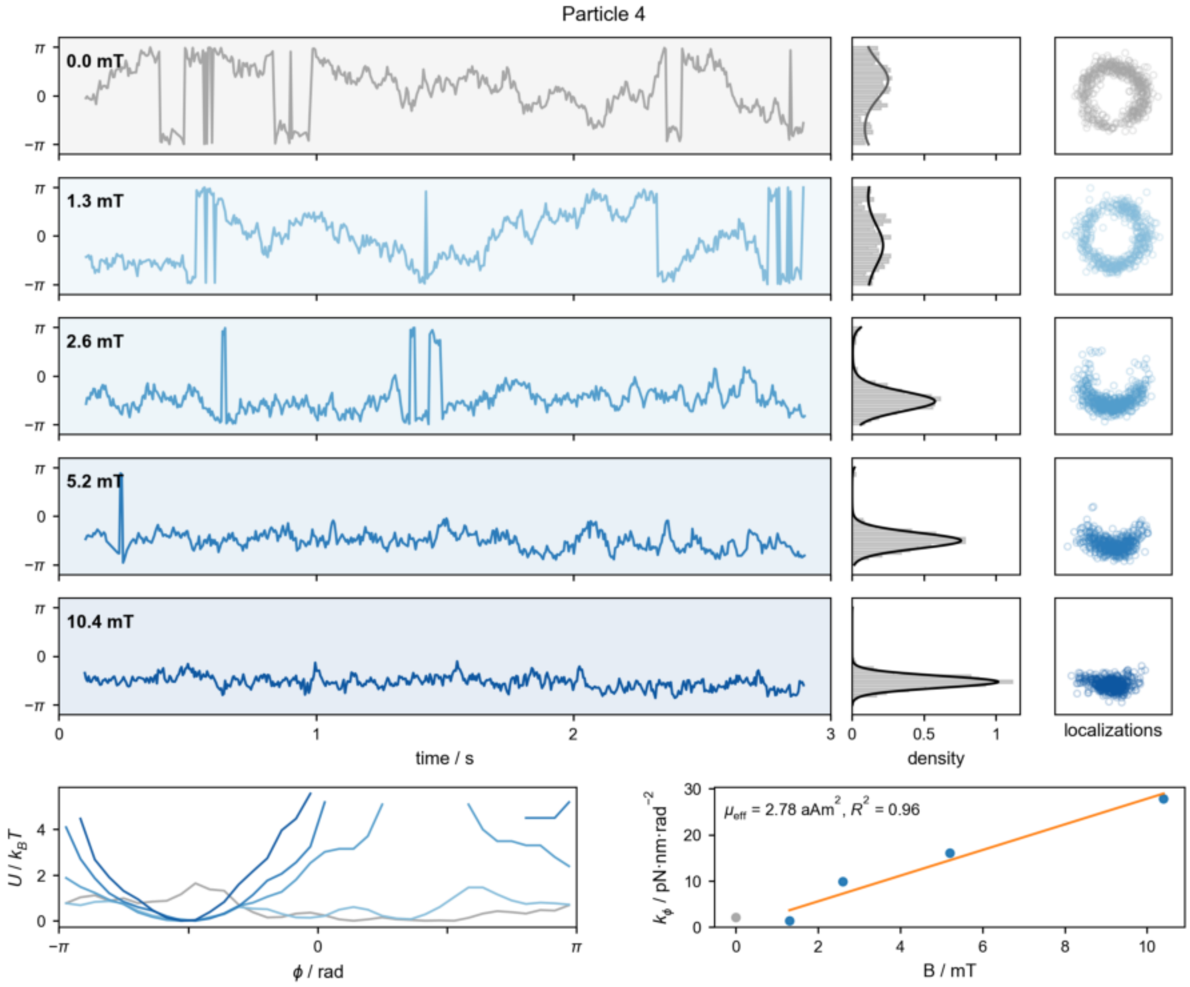


FIG. S17. **Exemplary 4-MNC MADONA rotor during clamping experiment**. Trajectories during clamping, angular localization histogram and van Mises fit, scatter plot of $x, y$- localizations. Energy landscapes (bottom left) and traps stiffness as function of the applied field (bottom right).

### 10. MONTE CARLO SIMULATIONS OF MAGNETIC PROPERTIES OF MADONAS

To capture the overal magnetic configuration of MADONAs in homogeneous magnetic fields and to estimate their net magnetization axis, magnetic moment and torques, we minimized the total magnetic energy. We used Metropolis Monte Carlo (MMC) energy minimization scheme for this purpose. The total magnetic energy $U_{\mathrm{i}}^{\mathrm{T}}$ of MNCs in external magnetic fields is given by a modified Stoner-Wohlfarth (SW) model for a single-domain MNC. The modfied SW model considers three energy terms: the Zeeman energy presenting particle-field interaction, the magnetic anisotropy energy for a cubic crystal (purely magnetocrystalline nature and ignoring $K_0$), and the magnetic dipole-dipole interaction energy, and is given by:

$$\frac{U_T}{k_B T} = -\xi_i \boldsymbol{m_i} \cdot \boldsymbol{B} + \frac{K_1 V_i}{k_B T}[\alpha_1^2\alpha_2^2 + \alpha_2^2\alpha_3^2 + \alpha_3^2\alpha_1^2] + (\frac{d_0}{r_{ij}})^3 \lambda \{\boldsymbol{m_i} \cdot \boldsymbol{m_j} - 3(\boldsymbol{m_i} \cdot \boldsymbol{r_{ij}})(\boldsymbol{m_j} \cdot \boldsymbol{r_{ij}})\}, \tag{16}$$

with

$$\xi_i = \frac{m_i B}{k_B T}, \tag{17}$$

and

$$\lambda = \frac{\mu_0 {m_0}^2}{4\pi k_B T {d_0}^3}, \tag{18}$$

in which $k_{\mathrm{B}}$ is the Boltzmann's constant in J K$^{-1}$, $T$ (295 k) is the temperature in K, $d_{\mathrm{i}}$ is the particle $i$ core diameter in nm, $r_{\mathrm{ij}} = \|r_{\mathrm{ij}}\|$ is the magnitude of the vector drawn from particles $i$ to $j$ in nm (Figure 4d), $\xi$ is the particle's dipole-field interaction parameter, $\lambda$ is a dimensionless dipole-dipole coupling parameter, $K_1$ is the first-order anisotropy coefficient in J m$^{-3}$, $m_0$ is the average magnetic moment, $d_0$ is the average particle core diameter. Items shown in bold are the unit vectors (e.g. $\boldsymbol{m_{\mathrm{i}}}$ is the magnetic moment unit vector of the particle $i$ and $m_{\mathrm{i}}= \|m_{\mathrm{i}}\|$). $\alpha_1 = \cos(\theta_1)$, $\alpha_2 = \cos(\theta_2)$, and $\alpha_3 = \cos(\theta_3)$ are the cosines of the angles between the magetization vector and the principle crystallographic axes of the crystal (Fig. S18a).

Our $Co_{0.4}Zn_{0.2}Fe_{2.4}O_4$ MNCs with a cubic crystal strcucture have a cubic magnetic anisotropy which is given by the second term im eq. 19 (Fig. S7a). The dominance of the cubic magnetic anisotropy can further be deduced by looking at the remanence to saturation magnetization ($M_{\mathrm{r}}/M_{\mathrm{s}}$) ratio of magnetic hysteresis loops at 5 K (Fig. S18b). The $M_{\mathrm{r}}/M_{\mathrm{s}}$ = 0.76 falls within the regime where the cubic magnetic anisotropy governs. Cobalt-doped zinc ferrite MNCs have the first-order magnetic anisotropy constant $K_1 > 0$, thus having six magnetic easy axes (Fig. S18a).

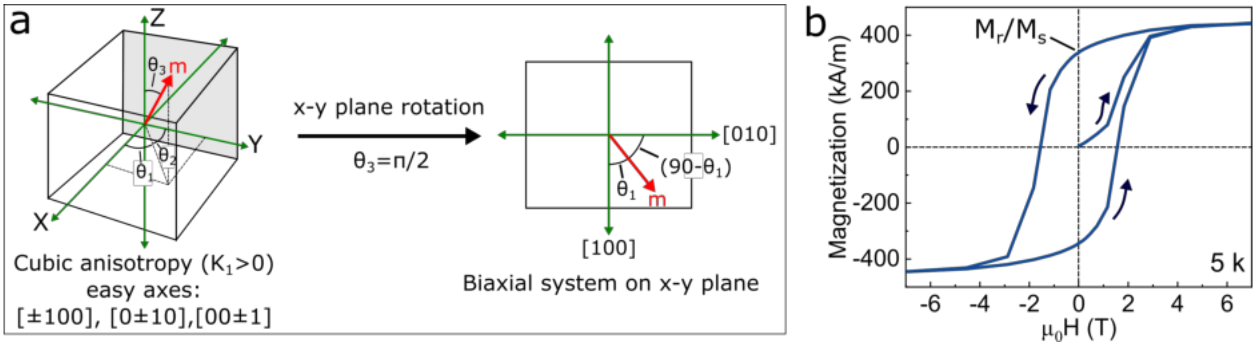


FIG. S18. **Magnetic anisotropy model of custom MNCs**.(a) Scheme of a single MNC with Cubic magnetic anisotropy and the first-order anisotropy constant $K_1$ with sixfold magnetization easy axes. Considering the magnetic rotation experiments in x-y plane, the symmetry from sixfold symmetry can be reduced to fourfold with two easy magnetization directions. This so-called biaxial system shows four energy minima at 0°, 90°, 180°, and 270°. (b) Magnetic hysteresis loops of individual MNCs measured at 5 K, showing the direction in which the magnetization loops are recorded.

We can simplify the anisotropy axis sytem from a sixfold to a fourfold symmetry with two magnetization easy directions: [100] and [010], since the magnetic actuation experiments are done on the x-y surface plane (i.e. $\theta_3 = 0$ and $\cos(\theta_3) = 0$) (Fig. S18a). By doing so, the cubic magnetic anisotropy energy term in eq. 19 will be simplified to:

$$\begin{aligned}\frac{E_a}{k_B T} &= \frac{K_1 V_i}{k_B T}(\alpha_1^2\alpha_2^2) = \frac{K_1 V_i}{k_B T}\cos^2(\theta_1)\cos^2(\theta_2)\\ \theta_2 &= \frac{\pi}{2} - \theta_1\\ \cos^2(\theta_2) &= sin^2(\theta_1)\\ \frac{E_a}{k_B T} &= \frac{K_1 V_i}{k_B T}\cos^2(\theta_1)\sin^2(\theta_1)\\ \frac{E_a}{k_B T} &= \frac{K_1 V_i}{4k_B T}\sin^2(2\theta_1)\end{aligned} \quad (19)$$

with

$$K_1 = 4K_{eff} \quad (20)$$

Through this simplification, the in x-y plane cubic magnetic anisotropy resembles the uniaxial anisotropy with four energy minima at 0°, 90°, 180°, and 270°.

The MMC simulations are 2D in-plane, in which the particle' magnetic moment is considered as a magnetic macrospin, pointing out from its center of coordination in a Cartesian x-y coordination system as shown in Fig. S18a.

At the beginning of each MMC simulation, the particle's magnetic moments and easy axes are randomly chosen in the x-y plane. Both vectors are set to be aligned to fulfill the magnetization in our ferrimagnetic MNCs with permanent dipole moment, where the magnetic moment is coupled to the easy axis. Next and at each MC sweep, a particle is chosen randomly and its magnetic moment and easy axis are rotated randomly for a maximum of 10° and 5°, respectively. The rotation of moment and easy axis vectors occurs in the same direction, with maximum 10° phase-lag between them. As the x-y coordinate will be set to the direction of the easy axis at the start of each MC simulation, the random rotation of the easy axis corresponds to the random physical rotation of MNCs (Figure 4d) on the 6HB. Due to ferrimagnetic nature of our MNCs, we considered that the alignment of the particle's moment with field requires a physical rotation of MNCs after being bound to the bundles (Figure 4d). As the MNCs are coupled to the 6HBs through three 8 nt double-stranded DNA, we hypothesize that the rotational freedom of bound MNCs is limited to a maximum of 90° toward the field. To give a full picture, we performed MMC simulations at 45°, 60°, and 90° rotational freedom. Next, at each new configuration, the total free energy is calculated and compared with the old configuration. If the total energy is reduced, the new configutaion is accepted. If the total energy is increased, the Metroplois criterion is applied and depending on the outcome, either the old or new configuration is accepted.

The following experimentally determined parameters were plugged into the MC simulations: $M_\mathrm{s}$ = 269 kA m$^{-1}$, $K_\mathrm{eff}$ = 50200 J m$^{-3}$. The particle size distribution was considerd to be log-normal with $\mu$ = 2.815 and $\sigma$ = 0.09, as obtained from TEM analyses (Figure 1). Each MC simulation is iterated for 500.000 MC sweeps to reach equilibrium. Uniform magnetic fields along the [0-10] direction with the field strength of 9 mT were used. From the MC simulations, we obtained equailibrium magnetic configuration, effective magnetic moment at R$_\mathrm{c-c}$ = 64 nm, and maximum magnetic torques. Our MC simulations capture the magnetic configuration of MADONAs in the clamping experiments.

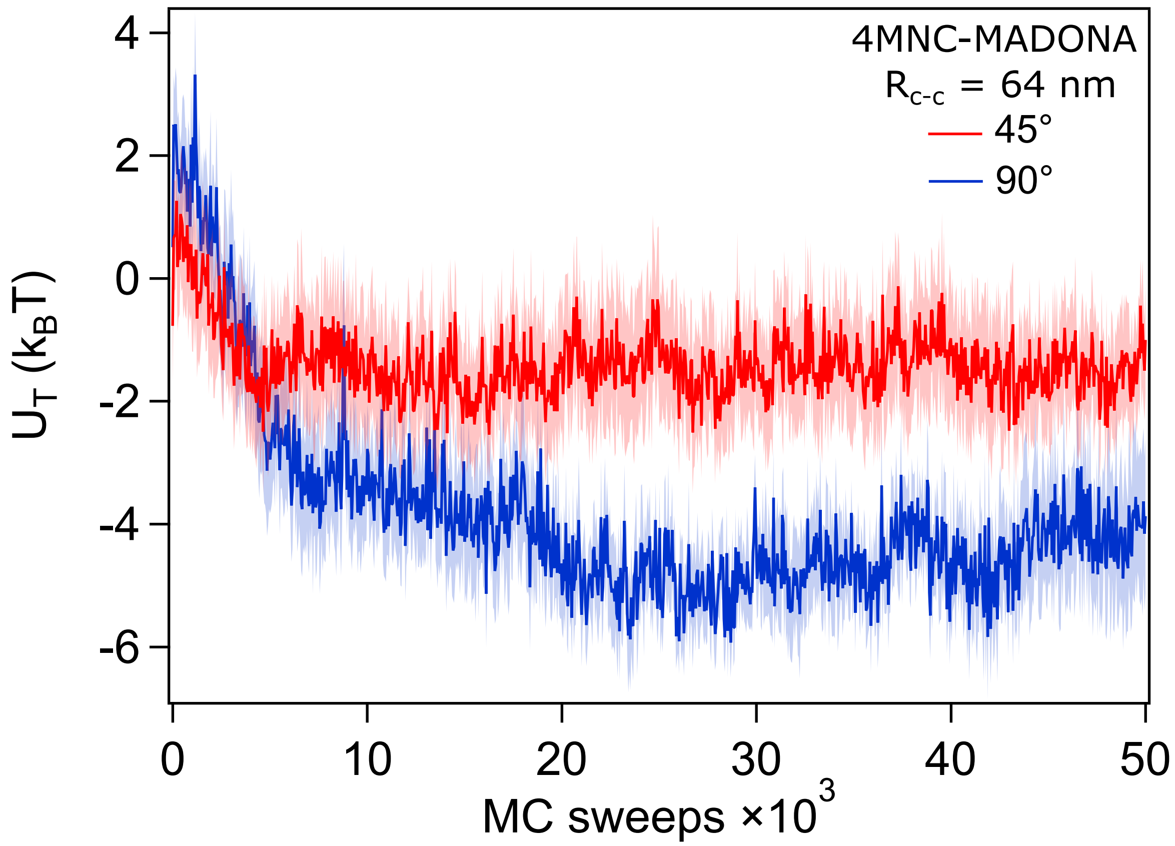


FIG. S19. **Total magnetic energy minimization using MMC simulations**. Changes in the total magnetic energy of 4-MNC MADONA at $R_{c-c}$ = 64 nm as a function of MMC sweeps at 45 and 90 ° rotational freedom. Data shown are the mean $\pm$ s.e.m of fifteen MC simulations at 9 mT magnetic fields. Errors are shown as shaded bands. Data are interpolated to 1000 points (cubic spline) for the sake of clarity and presentation. Only the first 50 k MMC sweeps out of the total 500 k are shown.

### 11. SCAFFOLD AND STAPLE SEQUENCES

**$7,249$ basepair scaffold version used for the 6hb structues:**

```
1         11        21        31        41        51        61        71
AAGCTTGGCACTGGCCGTCGTTTTACAACGTCGTGACTGGGAAAACCCTGGCGTTACCCAACTTAATCGCCTTGCAGCAC
81        91        101       111       121       131       141       151
ATCCCCCTTTCGCCAGCTGGCGTAATAGCGAAGAGGCCCGCACCGATCGCCCTTCCCAACAGTTGCGCAGCCTGAATGGC
161       171       181       191       201       211       221       231
GAATGGCGCTTTGCCTGGTTTCCGGCACCAGAAGCGGTGCCGGAAAGCTGGCTGGAGTGCGATCTTCCTGAGGCCGATAC
241       251       261       271       281       291       301       311
TGTCGTCGTCCCCTCAAACTGGCAGATGCACGGTTACGATGCGCCCATCTACACCAACGTGACCTATCCCATTACGGTCA
321       331       341       351       361       371       381       391
ATCCGCCGTTTGTTCCCACGGAGAATCCGACGGGTTGTTACTCGCTCACATTTAATGTTGATGAAAGCTGGCTACAGGAA
401       411       421       431       441       451       461       471
GGCCAGACGCGAATTATTTTTGATGGCGTTCCTATTGGTTAAAAAATGAGCTGATTTAACAAAAATTTAATGCGAATTTT
481       491       501       511       521       531       541       551
AACAAAATATTAACGTTTACAATTTAAATATTTGCTTATACAATCTTCCTGTTTTTGGGGCTTTTCTGATTATCAACCGG
561       571       581       591       601       611       621       631
GGTACATATGATTGACATGCTAGTTTTACGATTACCGTTCATCGATTCTCTTGTTTGCTCCAGACTCTCAGGCAATGACC
641       651       661       671       681       691       701       711
TGATAGCCTTTGTAGATCTCTCAAAAATAGCTACCCTCTCCGGCATTAATTTATCAGCTAGAACGGTTGAATATCATATT
721       731       741       751       761       771       781       791
GATGGTGATTTGACTGTCTCCGGCCTTTCTCACCCTTTTGAATCTTTACCTACACATTACTCAGGCATTGCATTTAAAAT
801       811       821       831       841       851       861       871
ATATGAGGGTTCTAAAAATTTTTATCCTTGCGTTGAAATAAAGGCTTCTCCCGCAAAAGTATTACAGGGTCATAATGTTT
881       891       901       911       921       931       941       951
TTGGTACAACCGATTTAGCTTTATGCTCTGAGGCTTTATTGCTTAATTTTGCTAATTCTTTGCCTTGCCTGTATGATTTA
961       971       981       991       1001      1011      1021      1031
TTGGATGTTAATGCTACTACTATTAGTAGAATTGATGCCACCTTTTCAGCTCGCGCCCCAAATGAAAATATAGCTAAACA
1041      1051      1061      1071      1081      1091      1101      1111
GGTTATTGACCATTTGCGAAATGTATCTAATGGTCAAACTAAATCTACTCGTTCGCAGAATTGGGAATCAACTGTTATAT
1121      1131      1141      1151      1161      1171      1181      1191
GGAATGAAACTTCCAGACACCGTACTTTAGTTGCATATTTAAAACATGTTGAGCTACAGCATTATATTCAGCAATTAAGC
1201      1211      1221      1231      1241      1251      1261      1271
TCTAAGCCATCCGCAAAAATGACCTCTTATCAAAAGGAGCAATTAAAGGTACTCTCTAATCCTGACCTGTTGGAGTTTGC
1281      1291      1301      1311      1321      1331      1341      1351
TTCCGGTCTGGTTCGCTTTGAAGCTCGAATTAAAACGCGATATTTGAAGTCTTTCGGGCTTCCTCTTAATCTTTTTGATG
1361      1371      1381      1391      1401      1411      1421      1431
CAATCCGCTTTGCTTCTGACTATAATAGTCAGGGTAAAGACCTGATTTTTGATTTATGGTCATTCTCGTTTTCTGAACTG
1441      1451      1461      1471      1481      1491      1501      1511
TTTAAAGCATTTGAGGGGGATTCAATGAATATTTATGACGATTCCGCAGTATTGGACGCTATCCAGTCTAAACATTTTAC
1521      1531      1541      1551      1561      1571      1581      1591
TATTACCCCCTCTGGCAAAACTTCTTTTGCAAAAGCCTCTCGCTATTTTGGTTTTTATCGTCGTCTGGTAAACGAGGGTT
1601      1611      1621      1631      1641      1651      1661      1671
ATGATAGTGTTGCTCTTACTATGCCTCGTAATTCCTTTTGGCGTTATGTATCTGCATTAGTTGAATGTGGTATTCCTAAA
1681      1691      1701      1711      1721      1731      1741      1751
TCTCAACTGATGAATCTTTCTACCTGTAATAATGTTGTTCCGTTAGTTCGTTTTATTAACGTAGATTTTTCTTCCCAACG
1761      1771      1781      1791      1801      1811      1821      1831
TCCTGACTGGTATAATGAGCCAGTTCTTAAAATCGCATAAGGTAATTCACAATGATTAAAGTTGAAATTAAACCATCTCA
1841      1851      1861      1871      1881      1891      1901      1911
AGCCCAATTTACTACTCGTTCTGGTGTTTCTCGTCAGGGCAAGCCTTATTCACTGAATGAGCAGCTTTGTTACGTTGATT
1921      1931      1941      1951      1961      1971      1981      1991
TGGGTAATGAATATCCGGTTCTTGTCAAGATTACTCTTGATGAAGGTCAGCCAGCCTATGCGCCTGGTCTGTACACCGTT
2001      2011      2021      2031      2041      2051      2061      2071
CATCTGTCCTCTTTCAAAGTTGGTCAGTTCGGTTCCCTTATGATTGACCGTCTGCGCCTCGTTCCGGCTAAGTAACATGG
```

```
2081      2091      2101      2111      2121      2131      2141      2151
AGCAGGTCGCGGATTTCGACACAATTTATCAGGCGATGATACAAATCTCCGTTGTACTTTGTTTCGCGCTTGGTATAATC
2161      2171      2181      2191      2201      2211      2221      2231
GCTGGGGGTCAAAGATGAGTGTTTTAGTGTATTCTTTTGCCTCTTTCGTTTTAGGTTGGTGCCTTCGTAGTGGCATTACG
2241      2251      2261      2271      2281      2291      2301      2311
TATTTTACCCGTTTAATGGAAACTTCCTCATGAAAAAGTCTTTAGTCCTCAAAGCCTCTGTAGCCGTTGCTACCCTCGTT
2321      2331      2341      2351      2361      2371      2381      2391
CCGATGCTGTCTTTCGCTGCTGAGGGTGACGATCCCGCAAAAGCGGCCTTTAACTCCCTGCAAGCCTCAGCGACCGAATA
2401      2411      2421      2431      2441      2451      2461      2471
TATCGGTTATGCGTGGGCGATGGTTGTTGTCATTGTCGGCGCAACTATCGGTATCAAGCTGTTTAAGAAATTCACCTCGA
2481      2491      2501      2511      2521      2531      2541      2551
AAGCAAGCTGATAAACCGATACAATTAAAGGCTCCTTTTGGAGCCTTTTTTTTGGAGATTTTCAACGTGAAAAAATTATT
2561      2571      2581      2591      2601      2611      2621      2631
ATTCGCAATTCCTTTAGTTGTTCCTTTCTATTCTCACTCCGCTGAAACTGTTGAAAGTTGTTTAGCAAAATCCCATACAG
2641      2651      2661      2671      2681      2691      2701      2711
AAAATTCATTTACTAACGTCTGGAAAGACGACAAAACTTTAGATCGTTACGCTAACTATGAGGGCTGTCTGTGGAATGCT
2721      2731      2741      2751      2761      2771      2781      2791
ACAGGCGTTGTAGTTTGTACTGGTGACGAAACTCAGTGTTACGGTACATGGGTTCCTATTGGGCTTGCTATCCCTGAAAA
2801      2811      2821      2831      2841      2851      2861      2871
TGAGGGTGGTGGCTCTGAGGGTGGCGGTTCTGAGGGTGGCGGTTCTGAGGGTGGCGGTACTAAACCTCCTGAGTACGGTG
2881      2891      2901      2911      2921      2931      2941      2951
ATACACCTATTCCGGGCTATACTTATATCAACCCTCTCGACGGCACTTATCCGCCTGGTACTGAGCAAAACCCCGCTAAT
2961      2971      2981      2991      3001      3011      3021      3031
CCTAATCCTTCTCTTGAGGAGTCTCAGCCTCTTAATACTTTCATGTTTCAGAATAATAGGTTCCGAAATAGGCAGGGGGC
3041      3051      3061      3071      3081      3091      3101      3111
ATTAACTGTTTATACGGGCACTGTTACTCAAGGCACTGACCCCGTTAAAACTTATTACCAGTACACTCCTGTATCATCAA
3121      3131      3141      3151      3161      3171      3181      3191
AAGCCATGTATGACGCTTACTGGAACGGTAAATTCAGAGACTGCGCTTTCCATTCTGGCTTTAATGAGGATTTATTTGTT
3201      3211      3221      3231      3241      3251      3261      3271
TGTGAATATCAAGGCCAATCGTCTGACCTGCCTCAACCTCCTGTCAATGCTGGCGGCGGCTCTGGTGGTGGTTCTGGTGG
3281      3291      3301      3311      3321      3331      3341      3351
CGGCTCTGAGGGTGGTGGCTCTGAGGGTGGCGGTTCTGAGGGTGGCGGCTCTGAGGGAGGCGGTTCCGGTGGTGGCTCTG
3361      3371      3381      3391      3401      3411      3421      3431
GTTCCGGTGATTTTGATTATGAAAAGATGGCAAACGCTAATAAGGGGGCTATGACCGAAAATGCCGATGAAAACGCGCTA
3441      3451      3461      3471      3481      3491      3501      3511
CAGTCTGACGCTAAAGGCAAACTTGATTCTGTCGCTACTGATTACGGTGCTGCTATCGATGGTTTCATTGGTGACGTTTC
3521      3531      3541      3551      3561      3571      3581      3591
CGGCCTTGCTAATGGTAATGGTGCTACTGGTGATTTTGCTGGCTCTAATTCCCAAATGGCTCAAGTCGGTGACGGTGATA
3601      3611      3621      3631      3641      3651      3661      3671
ATTCACCTTTAATGAATAATTTCCGTCAATATTTACCTTCCCTCCCTCAATCGGTTGAATGTCGCCCTTTTGTCTTTGGC
3681      3691      3701      3711      3721      3731      3741      3751
GCTGGTAAACCATATGAATTTTCTATTGATTGTGACAAAATAAACTTATTCCGTGGTGTCTTTGCGTTTCTTTTATATGT
3761      3771      3781      3791      3801      3811      3821      3831
TGCCACCTTTATGTATGTATTTTCTACGTTTGCTAACATACTGCGTAATAAGGAGTCTTAATCATGCCAGTTCTTTTGGG
3841      3851      3861      3871      3881      3891      3901      3911
TATTCCGTTATTATTGCGTTTCCTCGGTTTCCTTCTGGTAACTTTGTTCGGCTATCTGCTTACTTTTCTTAAAAAGGGCT
3921      3931      3941      3951      3961      3971      3981      3991
TCGGTAAGATAGCTATTGCTATTTCATTGTTTCTTGCTCTTATTATTGGGCTTAACTCAATTCTTGTGGGTTATCTCTCT
4001      4011      4021      4031      4041      4051      4061      4071
GATATTAGCGCTCAATTACCCTCTGACTTTGTTCAGGGTGTTCAGTTAATTCTCCCGTCTAATGCGCTTCCCTGTTTTTA
4081      4091      4101      4111      4121      4131      4141      4151
TGTTATTCTCTCTGTAAAGGCTGCTATTTTCATTTTTGACGTTAAACAAAAAATCGTTTCTTATTTGGATTGGGATAAAT
4161      4171      4181      4191      4201      4211      4221      4231
AATATGGCTGTTTATTTTGTAACTGGCAAATTAGGCTCTGGAAAGACGCTCGTTAGCGTTGGTAAGATTCAGGATAAAAT
4241      4251      4261      4271      4281      4291      4301      4311
TGTAGCTGGGTGCAAAATAGCAACTAATCTTGATTTAAGGCTTCAAAACCTCCCGCAAGTCGGGAGGTTCGCTAAAACGC
```

4321 4331 4341 4351 4361 4371 4381 4391

CTCGCGTTCTTAGAATACCGGATAAGCCTTCTATATCTGATTTGCTTGCTATTGGGCGCGGTAATGATTCCTACGATGAA

4401 4411 4421 4431 4441 4451 4461 4471

AATAAAAACGGCTTGCTTGTTCTCGATGAGTGCGGTACTTGGTTTAATACCCGTTCTTGGAATGATAAGGAAAGACAGCC

4481 4491 4501 4511 4521 4531 4541 4551

GATTATTGATTGGTTTCTACATGCTCGTAAATTAGGATGGGATATTATTTTTCTTGTTCAGGACTTATCTATTGTTGATA

4561 4571 4581 4591 4601 4611 4621 4631

AACAGGCGCGTTCTGCATTAGCTGAACATGTTGTTTATTGTCGTCGTCTGGACAGAATTACTTTACCTTTTGTCGGTACT

4641 4651 4661 4671 4681 4691 4701 4711

TTATATTCTCTTATTACTGGCTCGAAAATGCCTCTGCCTAAATTACATGTTGGCGTTGTTAAATATGGCGATTCTCAATT

4721 4731 4741 4751 4761 4771 4781 4791

AAGCCCTACTGTTGAGCGTTGGCTTTATACTGGTAAGAATTTGTATAACGCATATGATACTAAACAGGCTTTTTCTAGTA

4801 4811 4821 4831 4841 4851 4861 4871

ATTATGATTCCGGTGTTTATTCTTATTTAACGCCTTATTTATCACACGGTCGGTATTTCAAACCATTAAATTTAGGTCAG

4881 4891 4901 4911 4921 4931 4941 4951

AAGATGAAATTAACTAAAATATATTTGAAAAAGTTTTCTCGCGTTCTTTGTCTTGCGATTGGATTTGCATCAGCATTTAC

4961 4971 4981 4991 5001 5011 5021 5031

ATATAGTTATATAACCCAACCTAAGCCGGAGGTTAAAAAGGTAGTCTCTCAGACCTATGATTTTGATAAATTCACTATTG

5041 5051 5061 5071 5081 5091 5101 5111

ACTCTTCTCAGCGTCTTAATCTAAGCTATCGCTATGTTTTCAAGGATTCTAAGGGAAAATTAATTAATAGCGACGATTTA

5121 5131 5141 5151 5161 5171 5181 5191

CAGAAGCAAGGTTATTCACTCACATATATTGATTTATGTACTGTTTCCATTAAAAAAGGTAATTCAAATGAAATTGTTAA

5201 5211 5221 5231 5241 5251 5261 5271

ATGTAATTAATTTTGTTTTCTTGATGTTTGTTTCATCATCTTCTTTTGCTCAGGTAATTGAAATGAATAATTCGCCTCTG

5281 5291 5301 5311 5321 5331 5341 5351

CGCGATTTTGTAACTTGGTATTCAAAGCAATCAGGCGAATCCGTTATTGTTTCTCCCGATGTAAAAGGTACTGTTACTGT

5361 5371 5381 5391 5401 5411 5421 5431

ATATTCATCTGACGTTAAACCTGAAAATCTACGCAATTTCTTTATTTCTGTTTTACGTGCAAATAATTTTGATATGGTAG

5441 5451 5461 5471 5481 5491 5501 5511

GTTCTAACCCTTCCATTATTCAGAAGTATAATCCAAACAATCAGGATTATATTGATGAATTGCCATCATCTGATAATCAG

5521 5531 5541 5551 5561 5571 5581 5591

GAATATGATGATAATTCCGCTCCTTCTGGTGGTTTCTTTGTTCCGCAAAATGATAATGTTACTCAAACTTTTAAAATTAA

5601 5611 5621 5631 5641 5651 5661 5671

TAACGTTCGGGCAAAGGATTTAATACGAGTTGTCGAATTGTTTGTAAAGTCTAATACTTCTAAATCCTCAAATGTATTAT

5681 5691 5701 5711 5721 5731 5741 5751

CTATTGACGGCTCTAATCTATTAGTTGTTAGTGCTCCTAAAGATATTTTAGATAACCTTCCTCAATTCCTTTCAACTGTT

5761 5771 5781 5791 5801 5811 5821 5831

GATTTGCCAACTGACCAGATATTGATTGAGGGTTTGATATTTGAGGTTCAGCAAGGTGATGCTTTAGATTTTTCATTTGC

5841 5851 5861 5871 5881 5891 5901 5911

TGCTGGCTCTCAGCGTGGCACTGTTGCAGGCGGTGTTAATACTGACCGCCTCACCTCTGTTTTATCTTCTGCTGGTGGTT

5921 5931 5941 5951 5961 5971 5981 5991

CGTTCGGTATTTTTAATGGCGATGTTTTAGGGCTATCAGTTCGCGCATTAAAGACTAATAGCCATTCAAAAATATTGTCT

6001 6011 6021 6031 6041 6051 6061 6071

GTGCCACGTATTCTTACGCTTTCAGGTCAGAAGGGTTCTATCTCTGTTGGCCAGAATGTCCCTTTTATTACTGGTCGTGT

6081 6091 6101 6111 6121 6131 6141 6151

GACTGGTGAATCTGCCAATGTAAATAATCCATTTCAGACGATTGAGCGTCAAAATGTAGGTATTTCCATGAGCGTTTTTC

6161 6171 6181 6191 6201 6211 6221 6231

CTGTTGCAATGGCTGGCGGTAATATTGTTCTGGATATTACCAGCAAGGCCGATAGTTTGAGTTCTTCTACTCAGGCAAGT

6241 6251 6261 6271 6281 6291 6301 6311

GATGTTATTACTAATCAAAGAAGTATTGCTACAACGGTTAATTTGCGTGATGGACAGACTCTTTTACTCGGTGGCCTCAC

6321 6331 6341 6351 6361 6371 6381 6391

TGATTATAAAAACACTTCTCAGGATTCTGGCGTACCGTTCCTGTCTAAAATCCCTTTAATCGGCCTCCTGTTTAGCTCCC

6401 6411 6421 6431 6441 6451 6461 6471

GCTCTGATTCTAACGAGGAAAGCACGTTATACGTGCTCGTCAAAGCAACCATAGTACGCGCCCTGTAGCGGCGCATTAAG

6481 6491 6501 6511 6521 6531 6541 6551

CGCGGCGGGTGTGGTGGTTACGCGCAGCGTGACCGCTACACTTGCCAGCGCCCTAGCGCCCGCTCCTTTCGCTTTCTTCC

```
6561      6571      6581      6591      6601      6611      6621      6631
CTTCCTTTCTCGCCACGTTCGCCGGCTTTCCCCGTCAAGCTCTAAATCGGGGGCTCCCTTTAGGGTTCCGATTTAGTGCT
6641      6651      6661      6671      6681      6691      6701      6711
TTACGGCACCTCGACCCCAAAAAACTTGATTTGGGTGATGGTTCACGTAGTGGGCCATCGCCCTGATAGACGGTTTTTCG
6721      6731      6741      6751      6761      6771      6781      6791
CCCTTTGACGTTGGAGTCCACGTTCTTTAATAGTGGACTCTTGTTCCAAACTGGAACAACACTCAACCCTATCTCGGGCT
6801      6811      6821      6831      6841      6851      6861      6871
ATTCTTTTGATTTATAAGGGATTTTGCCGATTTCGGAACCACCATCAAACAGGATTTTCGCCTGCTGGGGCAAACCAGCG
6881      6891      6901      6911      6921      6931      6941      6951
TGGACCGCTTGCTGCAACTCTCTCAGGGCCAGGCGGTGAAGGGCAATCAGCTGTTGCCCGTCTCACTGGTGAAAAGAAAA
6961      6971      6981      6991      7001      7011      7021      7031
ACCACCCTGGCGCCCAATACGCAAACCGCCTCTCCCCGCGCGTTGGCCGATTCATTAATGCAGCTGGCACGACAGGTTTC
7041      7051      7061      7071      7081      7091      7101      7111
CCGACTGGAAAGCGGGCAGTGAGCGCAACGCAATTAATGTGAGTTAGCTCACTCATTAGGCACCCCAGGCTTTACACTTT
7121      7131      7141      7151      7161      7171      7181      7191
ATGCTTCCGGCTCGTATGTTGTGTGGAATTGTGAGCGGATAACAATTTCACACAGGAAACAGCTATGACCATGATTACGA
7201      7211      7221      7231      7241
ATTCGAGCTCGGTACCCGGGGATCCTCTAGAGTCGACCTGCAGGCATGC
```

**Staple sequences of the basic 6hb DNA origami nanostructure**

Staple sequences of the 6HB base structure used as MNC binding platform. Staple sequences are denoted in the form Name, starting helix [base position]end helix[base position], DNA sequence.

TABLE I: Extension sequences

| Type | Name | Sequence |
|---|---|---|
| basic 6HB | 1[154]3[167] | GCAACGGCGCCCACGCCTTGAGTAACAG |
| basic 6HB | 0[832]3[825] | AAGCAAACGAGCCGAAACGCAATAATAAAGAATAA |
| basic 6HB | 1[553]3[566] | AACAAAGAACAAACGAATCCTTGAAAAC |
| basic 6HB | 1[1036]3[1049] | CTTTGATAGCACTACGACGACAATAAAC |
| basic 6HB | 0[643]3[636] | TATAATGCTGCAAGCAAAAGGGCGACATCTGAGAG |
| basic 6HB | 1[280]3[293] | AAACACTTTGAAAATTGCTCAGTACCAG |
| basic 6HB | 1[343]3[356] | TATCATCGTTTCAGTGTATCACCGTACT |
| basic 6HB | 1[1015]3[1028] | TTAACCGAGAGCCGGGTAAAGTAATTCT |
| basic 6HB | 0[412]3[405] | CATACAGGATAGGTGTAGCGCGTTTTCAAACCGCC |
| basic 6HB | 0[286]3[279] | AAGGATATAGGAACAACCGCCACCCTCACGGGGTT |
| basic 6HB | 0[622]3[615] | GTTTTAAAGCTGGCATTGAGGGAGGGAAGAATTTA |
| basic 6HB | 1[322]3[335] | ACAAAGTAAAGGAAATAAGTATAGCCCG |
| basic 6HB | 1[448]3[461] | TGAAAGACACAGACAATAGGAACCCATG |
| basic 6HB | 0[370]3[363] | CCTCAGATCGGATTTAGCGTTTGCCATCCAGGAGG |
| basic 6HB | 0[580]3[573] | TCCATATGTTGGGAAAGGTGAATTATCAATAGCGA |
| basic 6HB | 1[133]3[146] | GACAGCATTCGGTCATAAGTTTTAACGG |
| basic 6HB | 1[511]3[524] | TAATCTTCATTTGACCTTGCTTCTGTAA |
| basic 6HB | 1[532]3[545] | ATTCATTATTAATTCTATTAATTAATTT |
| basic 6HB | 1[427]3[440] | GGGAACCGCGTAACATTTTCAGGGATAG |
| basic 6HB | 0[244]3[237] | TGCCTGATTCGCATAACCACCACCAGAGTATTAAG |
| basic 6HB | 1[1099]3[1112] | ATATCCACTGGTCATCCTGAACAAGAAA |
| basic 6HB | 1[889]3[902] | AAAACCATTGCGGACCAGTATAAAGCCA |
| basic 6HB | 1[700]3[713] | AGGACGTTTTAACGCCAATCGCAAGACA |
| basic 6HB | 0[748]3[741] | CCAGACCCGGGTACCATACATAAAGGTGCAAATAT |
| basic 6HB | 1[196]3[209] | AGGAAGTCAGCTTGCCTATTTCGGAACC |
| basic 6HB | 1[175]3[188] | AGGACTAGCCGACAATAAACAGTTAATG |
| basic 6HB | 0[538]3[531] | TAGTTTGGAAACCATAGAGCCAGCAAAAATCGTCG |
| basic 6HB | 0[979]3[972] | TACGCCAGGTTTGCCTAATATCAGAGAGCAGAGGC |
| basic 6HB | 0[874]3[867] | ATAAATCGTGAGCTGTAAGCAGATAGCCAGTATCA |

Continued on next page

| Type | Name | Sequence |
|---|---|---|
| basic 6HB | 0[790]3[783] | AAGACTTCCTGTGTAAGACTCCTTATTATGGTTTG |
| basic 6HB | 1[385]3[398] | CATGTTAAATGAATACCGCCACCCTCAG |
| basic 6HB | 0[391]3[384] | TAGCAAAACGGCGGTTTCGGTCATAGCCCCTCAGA |
| basic 6HB | 0[223]3[216] | AAGATTCAACGTTACAGCATTGACAGGATATTATT |
| basic 6HB | 0[916]3[909] | CCCCCTCTTCCAGTGAAATAGCAATAGCACGCTCA |
| basic 6HB | 0[853]3[846] | TACCCTGAAAGCCTGTTACCAGAAGGAAATTACTA |
| basic 6HB | 1[1057]3[1070] | CTTGCCTGTTATCTTCAGCTAATGCAGA |
| basic 6HB | 1[259]3[272] | AAGAGGCGCTCCAAAGGATTAGGATTAG |
| basic 6HB | 0[181]3[174] | CATCAATCCCAAAAGATATTCACAAACATGCCCGT |
| basic 6HB | 0[265]3[258] | CCTCATAATCAGCTCACCCTCAGAGCCGCAAGAGA |
| basic 6HB | 0[1063]3[1056] | GAGCACGAGAGAGTGAGAGAATAACATAAACATGT |
| basic 6HB | 1[574]3[587] | AATAAGGCTGAGCATAGCTTAGATTAAG |
| basic 6HB | 1[595]3[608] | AAACACCCGAATTAGAAGAGTCAATAGT |
| basic 6HB | 0[328]3[321] | GACCCTGGCCAGCTAGCCACCACCGGAAGGTTGAT |
| basic 6HB | 0[349]3[342] | CGGTTGTATGTGAGAATCAAAATCACCGGAATAGG |
| basic 6HB | 1[973]3[986] | TCAGTGAAGACTTTATTTTCGAGCCAGT |
| basic 6HB | 0[1105]3[1098] | CTTAATGGCGAAAAATTTTTTGTTTAACAGATAAG |
| basic 6HB | 1[301]3[314] | AGCGATTGCGAATAAGTGCCGTCGAGAG |
| basic 6HB | 1[826]3[839] | AAAGGAATGGCAATACACCGGAATCATA |
| basic 6HB | 1[616]3[629] | ATTGGGCAGTTACATCAAAATCATAGGT |
| basic 6HB | 0[664]3[657] | GCTTAGATAACGCCATGGTTTACCAGCGCTCCGGC |
| basic 6HB | 1[637]3[650] | TTTCAACGCCTGATACTACCTTTTTAAC |
| basic 6HB | 1[994]3[1007] | AGAGTCTATTTGAGGAATATAAAGTACC |
| basic 6HB | 0[475]3[468] | GGGCGCGCGACAGTAACCATCGATAGCATACCGTA |
| basic 6HB | 1[217]3[230] | TAAAATATTTCGAGCTGAAACATGAAAG |
| basic 6HB | 1[1078]3[1091] | AAACTATAGTTGAATGTTTATCAACAAT |
| basic 6HB | 0[1042]3[1035] | CTCGTTAGCCCTTCGGAAGCGCATTAGAGTCCAGA |
| basic 6HB | 1[931]3[944] | AGAGGGGCCGAACGTCGCCATATTTAAC |
| basic 6HB | 0[433]3[426] | TAGCATTTGGGCGCGTTTGCCTTTAGCGCACCCTC |
| basic 6HB | 0[1000]3[993] | AGGGATTGGTGGTTAAGTCAGAGGGTAAAATAAGA |
| basic 6HB | 0[727]3[720] | ACAGGTCAGGTCGAATAAAAGAAACGCAAAGAACG |
| basic 6HB | 1[679]3[692] | AACTGGCAGTAACATATGTAAATGCTGA |
| basic 6HB | 0[811]3[804] | AAAGATTCTCACAACCCAAAAGAACTGGTGATAAA |
| basic 6HB | 0[307]3[300] | GAGAAGCTTCGCGTCCTCAGAGCCGCCAGCGGATA |
| basic 6HB | 1[364]3[377] | GTCGAAATTTGCTATTTAGTACCGCCAC |
| basic 6HB | 1[742]3[755] | CGGAACATGCACGTATTTTAGTTAATTT |
| basic 6HB | 0[1084]3[1077] | GCGTACTCACGCTGATGAAAATAGCAGCACGCGCC |
| basic 6HB | 0[958]3[951] | TAGCGTCGGCCAACACAAGAATTGAGTTAACGCCA |
| basic 6HB | 0[937]3[930] | CGTCATATGCCAGCATAATAAGAGCAAGTTGAGAA |
| basic 6HB | 1[805]3[818] | TAATGCACTGATTGTAAGGCGTTAAATA |
| basic 6HB | 1[910]3[923] | TTTTGCAAGTTTGAACAGTAGGGCTTAA |
| basic 6HB | 0[706]3[699] | CCTTTAACCAGTGCCACGGAATAAGTTTTGCAAAT |
| basic 6HB | 0[160]3[153] | TTCTAGCCCGGTTGCCTCATTAAAGCCAGGTCAGT |
| basic 6HB | 1[406]3[419] | GGCGCAGTCGTCTTACCCTCAGAGCCAC |
| basic 6HB | 0[1021]3[1014] | GCTAAACTGAGACGATTAACTGAACACCGACAAAA |
| basic 6HB | 1[658]3[671] | ATTACCTGGGAGAATTAGGTTGGGTTAT |
| basic 6HB | 1[469]3[503] | TGTACAGACTACAAACACTGAGTTTCGTCAATAAATCAATATATGTGAG |
| basic 6HB | 1[721]3[734] | TACGTTAGAAATTGCGAGAAAACTTTTT |
| basic 6HB | 1[868]3[881] | CTCGTTTAGAAGGATATGCGTTATACAA |
| basic 6HB | 0[517]3[510] | TTCGCAACCGGCACTAGCACCATTACCATGAATAA |
| basic 6HB | 0[601]3[594] | TACGGTGGGGCCTCATTGACGGAAATTAACGCTGA |
| basic 6HB | 0[685]3[678] | AAGAGGTCACGACGCACAATCAATAGAAATAACTA |
| basic 6HB | 0[769]3[762] | TAATTCGTAATCATTGTTAGCAAACGTACATCTTC |
| basic 6HB | 1[763]3[776] | AGAAAGAACCTACCTGACCTAAATTTAA |
| basic 6HB | 1[847]3[860] | AAGAGCAATATTCCGAAAAAGCCTGTTT |
| basic 6HB | 0[559]3[552] | AATTCTGCGCCATTCGACTTGAGCCATTTCCCTTA |
| basic 6HB | 0[454]3[447] | CATCAATTCTGCCAAATCAGTAGCGACACAAGCCC |
| basic 6HB | 0[202]3[195] | GGCCGGATAAGCAAGCAGGTCAGACGATCCCCCTG |

Continued on next page

| Type | Name | Sequence |
|---|---|---|
| basic 6HB | 1[784]3[797] | TTTAGGACTGAATAAAATACCGACCGTG |
| basic 6HB | 0[496]4[483] | GTTTAGCGATCGCAGGCCGGAAACGTCA |
| basic 6HB | 0[895]3[888] | GTTCAGAGCGTTGCCCGAAGCCCTTTTTATTCTTA |
| basic 6HB | 5[98]0[98] | AAACAAGAGAATCGTACAAAGGCTATCA |
| basic 6HB | 1[238]3[251] | GAAGGCATGTATCGAGGCTGAGACTCCT |
| basic 6HB | 2[1266]0[1253] | CCATTAAAAATACCAGATAGAAAATCGG |
| basic 6HB | 2[1287]0[1274] | ACTGATAGCCCTAAAAAGCGTGTCGAGG |
| basic 6HB | 2[153]0[140] | GCATAACCGATATATCGGAACTTAATGC |
| basic 6HB | 4[146]1[153] | GAATGGACAATCATATGTACCTGATAAAGAGGGTA |
| basic 6HB | 4[1238]1[1245] | TTAGTTGTCAAAGGGCGAAAAAAGGGAGATTCTGG |
| basic 6HB | 4[1259]1[1266] | TTTTGAAATCAGGGCGATGGCAAGCACTACCCTTC |
| basic 6HB | 4[1217]1[1224] | GCTACAATAAAGAACGTGGACCTTGACGACGACCA |
| basic 6HB | 4[167]1[174] | AATAAATATAATCAGAAAAGCATGATATGGCTTTG |
| basic 6HB | 2[1245]0[1232] | CCACCAGCAGAAGAAAGGGACCCCCCGA |
| basic 6HB | 2[174]0[161] | ATGACAACAACCATCTACAGATCAACCG |
| basic 6HB | 2[1224]0[1211] | GAGGTGAGGCGGTCCAGTCACGGGAAAG |
| basic 6HB | 2[1203]0[1190] | ACACCGCCTGCAACTATTTACGAGAAAG |
| basic 6HB | 4[1196]1[1203] | TTACCAATTTGGAACAAGAGTACGTGGCATTGGCA |
| basic 6HB | 2[951]0[938] | ATTAAATCCTTTGCGTAATAGGCGGAAT |
| basic 6HB | 4[1112]1[1119] | AGAAACGTCCTGTTTGATGGTCCACACCATTACCG |
| basic 6HB | 2[552]0[539] | ATCAAGAAAACAAAACCCAAAGTAGATT |
| basic 6HB | 4[944]1[951] | AAGCCCATGCATTAATGAATCCAATACTTAAAATG |
| basic 6HB | 4[629]1[636] | TCAACCGGAAAGGGGGATGTGCTGTAGCGGTTTAA |
| basic 6HB | 2[720]0[707] | CGTAGATTTTCAGGTGGGAAGGAGAGTA |
| basic 6HB | 4[587]1[594] | TTCATTAAGGGCGATCGGTGCTCTGGAATGACGAG |
| basic 6HB | 2[1140]0[1127] | TGCTGAACCTCAAATTGCAACGCTGCGC |
| basic 6HB | 2[1056]0[1043] | AAAATATCTTTAGGTAGTAATTGCTTTC |
| basic 6HB | 4[713]1[720] | AAGACACCAAGCTTTGCCTGCAGGATTAAAAAATC |
| basic 6HB | 4[1049]1[1056] | AAAACAGACCGCCTGGCCCTGTATAACGAACATCA |
| basic 6HB | 4[1091]1[1098] | GTCAAAAGTTTGCCCCAGCAGCGCCGCTGCTGGTA |
| basic 6HB | 4[608]1[615] | GGTAAATTTCGCTATTACGCCATATGCAGTAGTAA |
| basic 6HB | 2[636]0[623] | TGCTTTGAATACCATTGAGATTCAACAT |
| basic 6HB | 2[1077]0[1064] | AGGAATTGAGGAAGGAGTAGACTTTGAC |
| basic 6HB | 4[797]1[804] | CATGATTGAAATTGTTATCCGAAGAGGAATTCAAC |
| basic 6HB | 2[762]0[749] | ATATCAAAATTATTACATTATAAGCGAA |
| basic 6HB | 4[860]1[867] | GAACAAAGGGGTGCCTAATGAAAAAATCCATAACC |
| basic 6HB | 4[671]1[678] | AATTCATAGGGTTTTCCCAGTCATTTTTTTTTAAG |
| basic 6HB | 4[818]1[825] | CGGAATATTCCACACAACATAGCGGATTAACGCCA |
| basic 6HB | 4[419]1[426] | TCAGACTCACGTTGGTGTAGAAACATCCATCATAA |
| basic 6HB | 2[405]0[392] | TCCAGACGTTAGTACTTAGCCAAAGAAT |
| basic 6HB | 4[440]1[447] | GAATCAAATCGTAACCGTGCATCTACTACCAACTT |
| basic 6HB | 2[594]0[581] | TTCATTTCAATTACCTTGCCCGTTTCAT |
| basic 6HB | 5[959]0[959] | GCGCGGGGAGAGGCGAATCCTGAGAAGA |
| basic 6HB | 1[490]0[476] | CTGGCTGTGGAAACAGTACCCAGTACAAACCAGGCTCATTTG |
| basic 6HB | 4[545]1[552] | TGGGAATGGCAAAGCGCCATTCGAACGATCAACGT |
| basic 6HB | 4[1007]1[1014] | CTGAACATTTCTTTTCACCAGAGGAGGCACGCAAA |
| basic 6HB | 2[888]0[875] | ACAAAGAAACCACCACCAGACAATGACC |
| basic 6HB | 4[923]1[930] | AAACAATCGGGAAACCTGTCGAATATTCTTTTGCC |
| basic 6HB | 4[398]1[405] | TCGGCATATTGACCGTAATGGGCAAGGCGGAACGA |
| basic 6HB | 4[902]1[909] | TATCTTAGCTCACTGCCCGCTAAATGCTGAGAGGC |
| basic 6HB | 2[804]0[791] | TTTGGATTATACTTATACCACAGCCCGA |
| basic 6HB | 4[1175]1[1182] | TTTCCAGTAGGGTTGAGTGTTAGAAAGCCGTCTGA |
| basic 6HB | 2[1161]0[1148] | GAAAAATCTAAAGCTGGAAATGCTGGCA |
| basic 6HB | 2[930]0[917] | TTATTAATTTTAAAAAAGAAGATTGAAT |
| basic 6HB | 2[573]0[560] | AAAGAAGATGATGACTGCTCAGATTCCC |
| basic 6HB | 2[783]0[770] | ATGGAAGGGTTAGATTCATCACGCGTTT |
| basic 6HB | 2[867]0[854] | GCGGAATTATCATCACACTATAGGTCTT |
| basic 6HB | 2[426]0[413] | GATCTAAAGTTTTGACGGTCAAATAAAT |

Continued on next page

| Type | Name | Sequence |
|---|---|---|
| basic 6HB | 2[468]0[455] | CGCCTGTAGCATTCGGACAGAAAGGTGG |
| basic 6HB | 4[524]1[531] | TCACCAGCGCTTCTGGTGCCGACCATTAACCGGAT |
| basic 6HB | 2[741]0[728] | AAAACAGAAATAAAATAAAACAACTCCA |
| basic 6HB | 4[566]1[573] | CCGTCACCAGGCTGCGCAACTAACAGTTTTCAGTG |
| basic 6HB | 4[734]1[741] | GCAACATCTCTAGAGGATCCCGGAAGCAGAACTAA |
| basic 6HB | 2[531]0[518] | ACATTTAACAATTTGACAAGAGATACAT |
| basic 6HB | 2[615]0[602] | AAATCGCGCAGAGGAGAACGAACTAAAG |
| basic 6HB | 2[1182]0[1169] | CGCTGAGAGCCAGCGCTCAATGAAAGGA |
| basic 6HB | 4[776]1[783] | CGCAGTAGGTCATAGCTGTTTCAAATATGTTGAGA |
| basic 6HB | 4[461]1[468] | GCACCGTGTTTGAGGGGACGAAGCTGAATGAACGG |
| basic 6HB | 2[846]0[833] | TGATTATCAGATGATTACGAGTAGTCAG |
| basic 6HB | 2[972]4[959] | ACAAACAATTCGACAACTCGTACATGTAATTTAGGATAACCC |
| basic 6HB | 2[1014]0[1001] | TCAATAGATAATACGTCCATCCGATTAA |
| basic 6HB | 4[839]1[846] | ACCGAGGGAAGCATAAAGTGTACTATTAGCATAGT |
| basic 6HB | 4[1133]1[1140] | TTATTTAAAATCGGCAAAATCCGGTCACAGGAAAA |
| basic 6HB | 2[825]0[812] | TCATCAATATAATCGATACATGCATCAA |
| basic 6HB | 4[1154]1[1161] | GTTACAAAATCAAAAGAATAGCTAGGGCACCTACA |
| basic 6HB | 4[482]1[489] | CCAATGAATCGGCCTCAGGAATATATTTGCATAGG |
| basic 6HB | 2[678]0[665] | GTACCTTTTACATCTATGCGAGCGGATG |
| basic 6HB | 2[993]0[980] | GATTTAGAAGTATTGGCCACCGGAACGG |
| basic 6HB | 2[384]0[371] | TTTCTGTATGGGATTCCGCGAAATAAAG |
| basic 6HB | 2[363]0[350] | AACAACTTTCAACAGCCTGATGCTAAAT |
| basic 6HB | 2[447]0[434] | AGCCCTCATAGTTAGAACTGAATAGTAG |
| basic 6HB | 2[699]0[686] | TCAGATGAATATACTCATTATTTTTGAT |
| basic 6HB | 4[881]1[888] | AAGAAAAAACTCACATTAATTAAACGAGGACGATA |
| basic 6HB | 4[986]1[993] | TTGAGCGGTATTGGGCGCCAGTTAGACAGAGTAAA |
| basic 6HB | 4[650]1[657] | CCAAAGAGCGATTAAGTTGGGGCTTAATATTGTGA |
| basic 6HB | 4[356]1[363] | TTTTCATCGAGTAACAACCCGGCATAAAAAATTGT |
| basic 6HB | 1[952]1[972] | TTTAGACTGGTGTTTTTATAA |
| basic 6HB | 2[657]0[644] | ACAATAACGGATTCTTTAATCTGCTGAA |
| basic 6HB | 2[909]0[896] | GTAACATTATCATTAAATAGCTTAAACA |
| basic 6HB | 2[1119]0[1106] | CCCTCAATCAATATGAACAATCGCCGCG |
| basic 6HB | 4[377]1[384] | CCCTTATCTCCGTGGGAACAAATTAAGCCCTGCTC |
| basic 6HB | 4[1028]1[1035] | CGGGAGAGGCAACAGCTGATTGAATCAGAATACTT |
| basic 6HB | 4[503]1[510] | TTAGCAACTCCAGCCAGCTTTATGGTCATCAAGAG |
| basic 6HB | 2[510]0[497] | ATTACCTTTTTTAAACCTTCAATAACCT |
| basic 6HB | 4[755]1[762] | GAAAATACGAGCTCGAATTCGAGCTTCATACAGGT |
| basic 6HB | 4[1070]1[1077] | CTTTACATGCAGCAAGCGGTCATGGTTGAGAACTC |
| basic 6HB | 2[1098]0[1085] | GTTGGCAAATCAACCGGCCTTACAGGGC |
| basic 6HB | 4[692]1[699] | ATTTTGTTTGTAAAACGACGGTTGCTCCACCAGTC |
| basic 6HB | 2[1035]0[1022] | ACAACTAATAGATTTTGTAGCAGCGGGA |
| basic 6HB | 3[100]3[125] | CATGGCTTTTGATGATACAGGAGTGT |
| basic 6HB | 0[139]3[132] | CGGAGAGAGCATGTAAGCGCAGTCTCTGACTGGTA |
| basic 6HB | 0[118]4[112] | AGAGATCATGAACGCGTTCCA |
| basic 6HB | 4[125]1[132] | AATTTACGTAATCGTAAAACTGGTAGCTAGCGAAA |
| basic 6HB | 2[132]2[115] | GCTGAGGCTTGCAGGGAG |
| basic 6HB | 1[115]0[119] | TCACCCTCAGCATTTTTG |
| marker binder 6HB | 1[1183]3[1196] | taagtgagacccgtacatatAATGGATAGTGCCACAAGAACGGGTATTtaagtgagacccgtacatat |
| marker binder 6HB | 0[1189]3[1182] | taagtgagacccgtacatatGAAGGGAGTTCCAGCGCTAACGAGCGTCATCATTCtaagtgagacccgtacatat |
| marker binder 6HB | 0[1168]3[1161] | taagtgagacccgtacatatGCGGGCGCCCGAGAAGCCTAATTTGCCAAATAATCtaagtgagacccgtacatat |
| marker binder 6HB | 0[1126]3[1119] | taagtgagacccgtacatatGTAACCAGGTTCCGTCCCAATCCAAATAAATAATAtaagtgagacccgtacatat |
| marker binder 6HB | 1[1120]3[1133] | taagtgagacccgtacatatCCAGCCATATCAAATCCCATCCTAATTTtaagtgagacccgtacatat |
| marker binder 6HB | 1[1162]3[1175] | taagtgagacccgtacatatTTTTGACAGCAAATGGCTGTCTTTCCTTtaagtgagacccgtacatat |
| marker binder 6HB | 0[1210]3[1203] | taagtgagacccgtacatatCCGGCGACCACTATTTTTATCCTGAATCAAACCAAtaagtgagacccgtacatat |
| marker binder 6HB | 1[1141]3[1154] | taagtgagacccgtacatatACGCTCAATCACCTTGTAGAAACCAATCtaagtgagacccgtacatat |
| marker binder 6HB | 0[1147]3[1140] | taagtgagacccgtacatatAGTGTAGCCTTATAAATAAACAGCCATAACGAGCAtaagtgagacccgtacatat |
| marker binder 6HB | 1[1267]3[1280] | taagtgagacccgtacatatTGACCTGAACATCGCCAATAGCAAGCAAtaagtgagacccgtacatat |
| marker binder 6HB | 1[1204]3[1217] | taagtgagacccgtacatatGATTCACAGTATTAGTACCGCACTCATCtaagtgagacccgtacatat |

Continued on next page

| Type | Name | Sequence |
|---|---|---|
| marker binder 6HB | 1[1246]3[1259] | taagtgagacccgtacatatCCAACAGGAACGAATCGTAGGAATCATTtaagtgagacccgtacatat |
| marker binder 6HB | 0[1273]3[1266] | taagtgagacccgtacatatTGCCGTACCACTACCGACTTGCGGGAGGACCGCGCtaagtgagacccgtacatat |
| marker binder 6HB | 1[1225]3[1238] | taagtgagacccgtacatatGTAATAATAAAACAAGCAAGCCGTTTTTtaagtgagacccgtacatat |
| marker binder 6HB | 0[1231]3[1224] | taagtgagacccgtacatatTTTAGAGTCCAACGCTATTTTGCACCCAGAGAACAtaagtgagacccgtacatat |
| marker binder 6HB | 0[1252]3[1245] | taagtgagacccgtacatatAACCCTAACCGTCTGCCTTAAATCAAGAATTTTCAtaagtgagacccgtacatat |
| marker binder 6HB | 1[1288]3[1310] | taagtgagacccgtacatatCGTGGCATGCGCGAATAGAAGGCTTATCCGGTATTTTtaagtgagacccgtacatat |
| marker binder 6HB | 0[1296]1[1287] | taagtgagacccgtacatatTTAAGTTTTTTGGGAAGAATAtaagtgagacccgtacatat |
| marker binder 6HB | 4[1310]3[1287] | taagtgagacccgtacatatTTTTCTAAGAACGCGAGGCGTTTTAGCGATCAGATtaagtgagacccgtacatat |
| marker binder 6HB | 4[1280]5[1296] | taagtgagacccgtacatatAACCTCCGTGAACCATCACCCAAATCTTtaagtgagacccgtacatat |
| marker binder 6HB | 2[1315]1[1315] | taagtgagacccgtacatatTTTGGCTATTAGTCTTTAACAGACAATATTTTTGAATTtaagtgagacccgtacatat |
| marker dye | - | ATTO655-TT-ATATGTACGGGTCTCACTTA |

**MNC binding staples Multi-MNC**

Staple sequences used for binding MNCs over the full length of the structure used for Multi-MNC design.

TABLE II: Multi-MNC binder staples

| Type | Name | Sequence |
|---|---|---|
| Multi-MNC binder | 4[1112]8[1112] | AGAAACGTCCTGTTTGATGGTCCACACCATTACCGAAAAAAAA |
| Multi-MNC binder | 4[566]8[566] | CCGTCACCAGGCTGCGCAACTAACAGTTTTCAGTGAAAAAAAA |
| Multi-MNC binder | 2[846]7[840] | TGATTATCAGATGATTACGAGTAGTCAGAAAAAAAA |
| Multi-MNC binder | 2[1014]7[1008] | TCAATAGATAATACGTCCATCCGATTAAAAAAAAAA |
| Multi-MNC binder | 2[1182]7[1176] | CGCTGAGAGCCAGCGCTCAATGAAAGGAAAAAAAAA |
| Multi-MNC binder | 2[363]7[357] | AACAACTTTCAACAGCCTGATGCTAAATAAAAAAAA |
| Multi-MNC binder | 2[216]7[210] | GTGAATTTCTTAAATTCCATTTGAGAAAAAAAAAAA |
| Multi-MNC binder | 2[1056]7[1050] | AAAATATCTTTAGGTAGTAATTGCTTTCAAAAAAAA |
| Multi-MNC binder | 4[293]8[293] | CCCTCAGGCCATCAAAAATAACTTTATTGACCCCCAAAAAAAA |
| Multi-MNC binder | 4[440]8[440] | GAATCAAATCGTAACCGTGCATCTACTACCAACTTAAAAAAAA |
| Multi-MNC binder | 1[952]8[965] | TTTAGACTGGTGTTTTTATAAAAAAAAAA |
| Multi-MNC binder | 4[1007]8[1007] | CTGAACATTTCTTTTCACCAGAGGAGGCACGCAAAAAAAAAAA |
| Multi-MNC binder | 4[251]8[251] | CCACCAGTAAATTTTTGTTAATATTTTAAAAACGAAAAAAAAA |
| Multi-MNC binder | 4[629]8[629] | TCAACCGGAAAGGGGGATGTGCTGTAGCGGTTTAAAAAAAAAA |
| Multi-MNC binder | 2[741]7[735] | AAAACAGAAATAAAATAAAACAACTCCAAAAAAAAA |
| Multi-MNC binder | 2[888]7[882] | ACAAAGAAACCACCACCAGACAATGACCAAAAAAAA |
| Multi-MNC binder | 4[356]8[356] | TTTTCATCGAGTAACAACCCGGCATAAAAAATTGTAAAAAAAA |
| Multi-MNC binder | 4[209]8[209] | GGTTGAGATATTTAAATTGTAAAAAGGGAAACGGGAAAAAAAA |
| Multi-MNC binder | 4[986]8[986] | TTGAGCGGTATTGGGCGCCAGTTAGACAGAGTAAAAAAAAAAA |
| Multi-MNC binder | 4[1070]8[1070] | CTTTACATGCAGCAAGCGGTCATGGTTGAGAACTCAAAAAAAA |
| Multi-MNC binder | 2[930]7[924] | TTATTAATTTTAAAAAAGAAGATTGAATAAAAAAAA |
| Multi-MNC binder | 2[1077]7[1071] | AGGAATTGAGGAAGGAGTAGACTTTGACAAAAAAAA |
| Multi-MNC binder | 4[1133]8[1133] | TTATTTAAAATCGGCAAAATCCGGTCACAGGAAAAAAAAAAAA |
| Multi-MNC binder | 2[804]7[798] | TTTGGATTATACTTATACCACAGCCCGAAAAAAAAA |
| Multi-MNC binder | 2[951]7[945] | ATTAAATCCTTTGCGTAATAGGCGGAATAAAAAAAA |
| Multi-MNC binder | 4[545]8[545] | TGGGAATGGCAAAGCGCCATTCGAACGATCAACGTAAAAAAAA |
| Multi-MNC binder | 5[959]7[966] | GCGCGGGGAGAGGCGAATCCTGAGAAGAAAAAAAAA |
| Multi-MNC binder | 2[699]7[693] | TCAGATGAATATACTCATTATTTTTGATAAAAAAAA |
| Multi-MNC binder | 2[279]7[273] | TCTCCAAAAAAAAGAAAAGAATTAGAACAAAAAAAA |
| Multi-MNC binder | 2[426]7[420] | GATCTAAAGTTTTGACGGTCAAATAAATAAAAAAAA |
| Multi-MNC binder | 2[636]7[630] | TGCTTTGAATACCATTGAGATTCAACATAAAAAAAA |
| Multi-MNC binder | 4[482]8[482] | CCAATGAATCGGCCTCAGGAATATATTTGCATAGGAAAAAAAA |
| Multi-MNC binder | 2[993]7[987] | GATTTAGAAGTATTGGCCACCGGAACGGAAAAAAAA |
| Multi-MNC binder | 4[797]8[797] | CATGATTGAAATTGTTATCCGAAGAGGAATTCAACAAAAAAAA |

Continued on next page

| Type | Name | Sequence |
|---|---|---|
| Multi-MNC binder | 2[762]7[756] | ATATCAAAATTATTACATTATAAGCGAAAAAAAAAA |
| Multi-MNC binder | 4[944]8[944] | AAGCCCATGCATTAATGAATCCAATACTTAAAATGAAAAAAAA |
| Multi-MNC binder | 2[783]7[777] | ATGGAAGGGTTAGATTCATCACGCGTTTAAAAAAAA |
| Multi-MNC binder | 2[195]7[189] | ATACCGATAGTTGCAAGACTTAAATCACAAAAAAAA |
| Multi-MNC binder | 2[1035]7[1029] | ACAACTAATAGATTTTGTAGCAGCGGGAAAAAAAAA |
| Multi-MNC binder | 4[671]8[671] | AATTCATAGGGTTTTCCCAGTCATTTTTTTTTAAGAAAAAAAA |
| Multi-MNC binder | 4[419]8[419] | TCAGACTCACGTTGGTGTAGAAACATCCATCATAAAAAAAAAA |
| Multi-MNC binder | 2[825]7[819] | TCATCAATATAATCGATACATGCATCAAAAAAAAAA |
| Multi-MNC binder | 2[1140]7[1134] | TGCTGAACCTCAAATTGCAACGCTGCGCAAAAAAAA |
| Multi-MNC binder | 4[335]8[335] | GAACCAGTTCATCAACATTAAACCAAAAAGATTTGAAAAAAAA |
| Multi-MNC binder | 2[594]7[588] | TTCATTTCAATTACCTTGCCCGTTTCATAAAAAAAA |
| Multi-MNC binder | 4[839]8[839] | ACCGAGGGAAGCATAAAGTGTACTATTAGCATAGTAAAAAAAA |
| Multi-MNC binder | 4[818]8[818] | CGGAATATTCCACACAACATAGCGGATTAACGCCAAAAAAAAA |
| Multi-MNC binder | 2[657]7[651] | ACAATAACGGATTCTTTAATCTGCTGAAAAAAAAAA |
| Multi-MNC binder | 2[1098]7[1092] | GTTGGCAAATCAACCGGCCTTACAGGGCAAAAAAAA |
| Multi-MNC binder | 2[1119]7[1113] | CCCTCAATCAATATGAACAATCGCCGCGAAAAAAAA |
| Multi-MNC binder | 4[902]8[902] | TATCTTAGCTCACTGCCCGCTAAATGCTGAGAGGCAAAAAAAA |
| Multi-MNC binder | 2[867]7[861] | GCGGAATTATCATCACACTATAGGTCTTAAAAAAAA |
| Multi-MNC binder | 2[447]7[441] | AGCCCTCATAGTTAGAACTGAATAGTAGAAAAAAAA |
| Multi-MNC binder | 2[342]7[336] | CGGAGTGAGAATAGACAACGGACATTATAAAAAAAA |
| Multi-MNC binder | 4[650]8[650] | CCAAAGAGCGATTAAGTTGGGGCTTAATATTGTGAAAAAAAAA |
| Multi-MNC binder | 2[510]7[504] | ATTACCTTTTTTAAACCTTCAATAACCTAAAAAAAA |
| Multi-MNC binder | 4[272]8[272] | GAGCCACCATTTTTTAACCAAAAAATTTTACACTAAAAAAAAA |
| Multi-MNC binder | 2[531]7[525] | ACATTTAACAATTTGACAAGAGATACATAAAAAAAA |
| Multi-MNC binder | 2[678]7[672] | GTACCTTTTACATCTATGCGAGCGGATGAAAAAAAA |
| Multi-MNC binder | 4[860]8[860] | GAACAAAGGGGTGCCTAATGAAAAAATCCATAACCAAAAAAAA |
| Multi-MNC binder | 2[909]7[903] | GTAACATTATCATTAAATAGCTTAAACAAAAAAAAA |
| Multi-MNC binder | 4[188]8[188] | TGGCCTTACAGGAAGATTGTAGACAGTCTTTCATGAAAAAAAA |
| Multi-MNC binder | 4[734]8[734] | GCAACATCAGTTGAGGATCCCGGAAGCAGAACTAAAAAAAAAA |
| Multi-MNC binder | 4[881]8[881] | AAGAAAAAACTCACATTAATTAAACGAGGACGATAAAAAAAAA |
| Multi-MNC binder | 4[1175]8[1175] | TTTCCAGTAGGGTTGAGTGTTAGAAAGCCGTCTGAAAAAAAAA |
| Multi-MNC binder | 4[461]8[461] | GCACCGTGTTTGAGGGGACGAAGCTGAATGAACGGAAAAAAAA |
| Multi-MNC binder | 2[552]7[546] | ATCAAGAAAACAAAACCCAAAGTAGATTAAAAAAAA |
| Multi-MNC binder | 4[503]8[503] | TTAGCAACTCCAGCCAGCTTTATGGTCATCAAGAGAAAAAAAA |
| Multi-MNC binder | 2[615]7[609] | AAATCGCGCAGAGGAGAACGAACTAAAGAAAAAAAA |
| Multi-MNC binder | 2[468]7[462] | CGCCTGTAGCATTCGGACAGAAAGGTGGAAAAAAAA |
| Multi-MNC binder | 4[524]8[524] | TCACCAGCGCTTCTGGTGCCGACCATTAACCGGATAAAAAAAA |
| Multi-MNC binder | 1[490]7[483] | CTGGCTGTGGAAACAGTACCCAGTACAAACCAGGCTCATTTGAAAAAAAA |
| Multi-MNC binder | 2[720]7[714] | CGTAGATTTTCAGGTGGGAAGGAGAGTAAAAAAAAA |
| Multi-MNC binder | 2[300]7[294] | ATAATTTTTTCACGCATCTTTTCAACGCAAAAAAAA |
| Multi-MNC binder | 4[923]8[923] | AAACAATCGGGAAACCTGTCGAATATTCTTTTGCCAAAAAAAA |
| Multi-MNC binder | 2[1161]7[1155] | GAAAAATCTAAAGCTGGAAATGCTGGCAAAAAAAAA |
| Multi-MNC binder | 4[314]8[314] | CCGCCTCCTGGCCTTCCTGTATAATACTGCGCGAAAAAAAAAA |
| Multi-MNC binder | 2[321]7[315] | CAACTAAAGGAATTATACCAATTTGCGGAAAAAAAA |
| Multi-MNC binder | 4[398]8[398] | TCGGCATATTGACCGTAATGGGCAAGGCGGAACGAAAAAAAAA |
| Multi-MNC binder | 4[1091]8[1091] | GTCAAAAGTTTGCCCCAGCAGCGCCGCTGCTGGTAAAAAAAAA |
| Multi-MNC binder | 2[384]7[378] | TTTCTGTATGGGATTCCGCGAAATAAAGAAAAAAAA |
| Multi-MNC binder | 2[237]7[231] | GTTTATCAGCTTGCCGTAATGGTAGGTAAAAAAAAA |
| Multi-MNC binder | 4[713]8[713] | AAGACACCAAGCTTCGTGAGCAGGATTAAAAAATCAAAAAAAA |
| Multi-MNC binder | 4[692]8[692] | ATTTTGTTTGTAAAACGACGGTTGCTCCACCAGTCAAAAAAAA |
| Multi-MNC binder | 4[587]8[587] | TTCATTAAGGGCGATCGGTGCTCTGGAATGACGAGAAAAAAAA |
| Multi-MNC binder | 4[1154]8[1154] | GTTACAAAATCAAAAGAATAGCTAGGGCACCTACAAAAAAAAA |
| Multi-MNC binder | 2[573]7[567] | AAAGAAGATGATGACTGCTCAGATTCCCAAAAAAAA |
| Multi-MNC binder | 4[1049]8[1049] | AAAACAGACCGCCTGGCCCTGTATAACGAACATCAAAAAAAAA |
| Multi-MNC binder | 4[776]8[776] | CGCAGTAGGTCATAGCTGTTTCAAATATGTTGAGAAAAAAAAA |
| Multi-MNC binder | 2[405]7[399] | TCCAGACGTTAGTACTTAGCCAAAGAATAAAAAAAA |

Continued on next page

| Type | Name | Sequence |
|---|---|---|
| Multi-MNC binder | 4[1028]8[1028] | CGGGAGAGGCAACAGCTGATTGAATCAGAATACTTAAAAAAAA |
| Multi-MNC binder | 4[755]8[755] | GAAAATACGAGCTCGAATTCGAGCTTCATACAGGTAAAAAAAA |
| Multi-MNC binder | 2[258]7[252] | AAGGAGCCTTTAATCCAACCTAATGCAAAAAAAAAA |
| Multi-MNC binder | 4[377]8[377] | CCCTTATCTCCGTGGAACAAATTAAGCCCTGCTCAAAAAAAA |
| Multi-MNC binder | 4[608]8[608] | GGTAAATTTCGCTATTACGCCATATGCAGTAGTAAAAAAAAAA |
| Multi-MNC binder | 4[230]8[230] | CCGCCGCATATTTTGTTAAAAGTAATGTCCACTACAAAAAAAA |

**MNC binding staples of the 4-MNC structure**

Staple sequences used for binding of MNCS on the 4-MNC structure.

TABLE III: 4 MNC-binding staples

| Type | Name | Sequence |
|---|---|---|
| MNC1 | 7[678]3[678] | AAAAAAAAAAGAGGTCACGACGCACAATCAATAGAAATAACTA |
| MNC1 | 7[699]3[703] | AAAAAAAACCTTTAACCAGTGCCACGGAATAAGTTTTGCAAATCCAA |
| MNC1 | 7[720]3[720] | AAAAAAAAACAGGTCAGGTCGAATAAAAGAAACGCAAAGAACG |
| MNC1 | 8[686]3[692] | AAAAAAAAAACTGGCAGTAACATATGTAAATGCTGA |
| MNC1 | 8[707]2[693] | AAAAAAAAAGGACGTTTTAACGTCAGATG |
| MNC1 | 8[728]3[734] | AAAAAAAATACGTTAGAAATTGCGAGAAAACTTTTT |
| MNC2 | 7[573]3[573] | AAAAAAAATCCATATGTTGGGAAAGGTGAATTATCAATAGCGA |
| MNC2 | 7[594]3[594] | AAAAAAAATACGGTGGGGCCTCATTGACGGAAATTAACGCTGA |
| MNC2 | 7[615]3[615] | AAAAAAAAGTTTTAAAGCTGGCATTGAGGGAGGGAAGAATTTA |
| MNC2 | 8[581]3[587] | AAAAAAAAAAATAAGGCTGAGCATAGCTTAGATTAAG |
| MNC2 | 8[602]3[608] | AAAAAAAAAAAACACCCGAATTAGAAGAGTCAATAGT |
| MNC2 | 8[623]3[629] | AAAAAAAAAATTGGGCAGTTACATCAAAATCATAGGT |
| MNC3 | 7[783]3[783] | AAAAAAAAAAGACTTCCTGTGTAAGACTCCTTATTATGGTTTG |
| MNC3 | 7[804]3[804] | AAAAAAAAAAAAGATTCTCACAACCCAAAAGAACTGGTGATAAA |
| MNC3 | 7[825]3[825] | AAAAAAAAAAGCAAACGAGCCGAAACGCAATAATAAAGAATAA |
| MNC3 | 8[791]3[797] | AAAAAAAATTTAGGACTGAATAAAATACCGACCGTG |
| MNC3 | 8[812]3[818] | AAAAAAAATAATGCACTGATTGTAAGGCGTTAAATA |
| MNC3 | 8[833]3[839] | AAAAAAAAAAAAGGAATGGCAATACACCGGAATCATA |
| MNC4 | 7[468]3[468] | AAAAAAAAGGGCGCGCGACAGTAACCATCGATAGCATACCGTA |
| MNC4 | 7[489]4[483] | AAAAAAAAGTTTAGCGATCGCAGGCCGGAAACGTCA |
| MNC4 | 7[510]3[510] | AAAAAAAATTCGCAACCGGCACTAGCACCATTACCATGAATAA |
| MNC4 | 8[476]3[503] | AAAAAAAATGTACAGACTACAAACACTGAGTTTCGTCAATAAATCAATATATGTGAG |
| MNC4 | 8[497]0[476] | AAAAAAAACTGGCTGTGGAAACAGTACCCAGTACAAACCAGGCTCATTTG |
| MNC4 | 8[518]3[524] | AAAAAAAATAATCTTCATTTGACCTTGCTTCTGTAA |

**Modifications staples**

Sequences of modification staples for either blunt end or passivated 6HB variant as well as for the addition of the two Biotin pivots.

TABLE IV: Modification staples

| Type | Name | Sequence |
|---|---|---|
| Blunt end 6HB | 0[118]4[100] | AGAGATCATGAACGCGTTCCAGTAAGCGTCATA |
| Blunt end 6HB | 2[132]2[103] | GCTGAGGCTTGCAGGGAGTTAAAGGCCGCT |
| Blunt end 6HB | 1[103]0[119] | TTTGCGGGATCGTCACCCTCAGCATTTTTG |

Continued on next page

| Type | Name | Sequence |
|---|---|---|
| Blunt end 6HB | 5[86]0[86] | AGAGTCTGGAGCAAACAAGAGAATCGTACAAAGGCTATCAGGTCATTGCCTG |
| Biotin end pivot | 1[196]3[209] | AGGAAGTCAGCTTGCCTATTTCGGAACCTT/3Bio/ |
| End passivation | 0[118]4[94] | AGAGATCATGAACGCGTTCCAGTAAGCGTCATATTTTTT |
| End passivation | 2[132]2[97] | GCTGAGGCTTGCAGGGAGTTAAAGGCCGCTTTTTTT |
| End passivation | 1[97]0[119] | TTTTTTTTTGCGGGATCGTCACCCTCAGCATTTTTG |
| End passivation | 5[80]0[80] | TTTTTTAGAGTCTGGAGCAAACAAGAGAATCGTACAAAGGCTATCAGGTCATTGCCTGTTTTTT |
| End passivation | 3[94]3[125] | TTTTTTCATGGCTTTTGATGATACAGGAGTGT |
| Mid pivot adapter | 5[714]1[720] | TGCCTGCAGGATTAAAAAATC |
| Mid pivot adapter | 2[692]0[686] | AATATACTCATTATTTTTGAT |
| Mid Biotin pivot | 3[704]5[713] | /5Biosg/TT TCGCAAGACAAAGACACCAAGCTT |
